# Stagewise identification of significantly mutated and dysregulated gene clusters in major molecular classes of breast invasive carcinoma

**DOI:** 10.1101/2020.05.30.125260

**Authors:** Shivangi Agarwal, Sanoj Naik, Pratima Kumari, Sandip K. Mishra, Amit K. Adhya, Sushil K. Kashaw, Anshuman Dixit

## Abstract

Breast invasive carcinoma (BRCA) is most malignant and leading cause of death in women. The efforts are ongoing for improvement in early detection, prevention and treatment. Therefore, identification of biomarkers/candidate genes has become very important. The current work includes comprehensive analysis of RNA-sequencing data of 1097 BRCA samples and 114 normal adjacent tissues to identify dysregulated genes in major molecular classes of BRCA in various clinical stages. Huge number of dysregulated genes were found, some were stage-specific, and others were common. The pathways as interferon signaling, tryptophan degradation III, granulocyte adhesion & diapedesis and catecholamine biosynthesis were found in ER/PR+/HER-2- (p-value<0.010), pathways as RAR activation, adipogenesis, role of JAK1, JAK2 in interferon signaling, TGF-β and STAT3 signaling (p-value<0.014) intricated in ER/PR-/HER-2+ and pathways as IL-1/IL-8 signaling, TNFR1/TNFR2 signaling, TWEAK and relaxin signaling (p-value<0.005) were found in triple negative breast cancer. The genes were clustered based on their mutation profile which revealed nine mutated clusters, some of which were known to be involved in well-known cancer signalling pathways while others were less characterized. Each cluster was analyzed in detail which led us to discover genes viz. NLGN3, MAML2, TTN, SYNE1, ANK2 as candidates in BRCA. The genes were found to be involved in important processes as chemotaxis, axon guidance, notch binding, cell-adhesion-molecule binding etc. They are central genes in the protein-protein-interaction network indicating they can have important regulatory roles. The qRT-PCR and western blot confirmed our findings in breast cancer cell-lines. Further, immunohistochemistry corroborated the results in ~100 tissue samples. The genes can be used as biomarker in BRCA.

## Introduction

BRCA is the most common cancer in females. As per Globocan 2018, around 2,088,849 new breast cancer cases were reported in 2018. The late detection and emergence of resistance to existing therapy pose a significant challenge in the management of BRCA [1]. Moreover, in-spite of recent advances in the understanding of the disease and development of new therapies and preventive measures, the five-year survival rate is still unsatisfactory [2, 3]. Therefore, new targets are required for early detection and treatment to improve overall survival.

The diverse cellular processes are carried out by small molecular machines formed of a myriad of protein complexes. An understanding of these molecular complexes can shed an important light on their functions and the processes they are involved into. The analysis of PPIs supplemented by vast amount of data available for gene expression, mutations, pathway and ontologies can lead to greater understanding and insights into the molecular mechanisms underlying the pathology of carcinogenesis [4].

Therefore, in the present work, an in-depth analysis of RNA-sequencing data obtained from the cancer genome atlas (TCGA) for breast invasive carcinoma is reported. The dysregulated genes in three molecular classes of BRCA (ER/PR+/HER-2-; ER/PR-/HER-2+; triple negative) in early and late stage have been identified. Further, different clusters of genes in various classes were investigated for their involvement in the biological processes. The subnetworks were found to contain frequently mutated genes in BRCA patients which can serve as therapeutic targets as well as biomarkers. The identified genes were further analyzed for their mutational patterns and effect on overall survival of patients. The findings were finally validated by qRT-PCR, western-blot in cell lines and immunohistochemistry using ~100 tissue samples.

## Results

### Differential expression analysis

The differential expression analysis was done separately for each of the five classes viz. (i) Early stage ER/PR+/HER-2-(ii) Late stage ER/PR+/HER-2-(iii) Early stage ER/PR-/HER-2+ (iv) Early stage triple negative breast cancer (TNBC) and (v) Late stage TNBC using tumor and normal adjacent tissue (NAT) samples. The number of upregulated and downregulated genes in each class is reported in table 2 and supplementary tables S1a-e. There were many dysregulated genes which were common among classes (Fig. 2a, 2b, supplementary table S2a) whereas some were class and stage specific (supplementary table S2b). Interestingly, some of the genes had inverse expression i.e. were upregulated in one class and downregulated in other (table 3, Fig. 2c, 2d, supplementary table S2c). These observations indicate some common and stage specific mechanisms underlie the pathology of different molecular classes of BRCA (Fig. 3a). The biological pathways were analyzed in each class (Fig. 3b, supplementary table S3 a, S3b, S3c, S3d, S3e) and significant pathways have been discussed.

**Figure 1.**
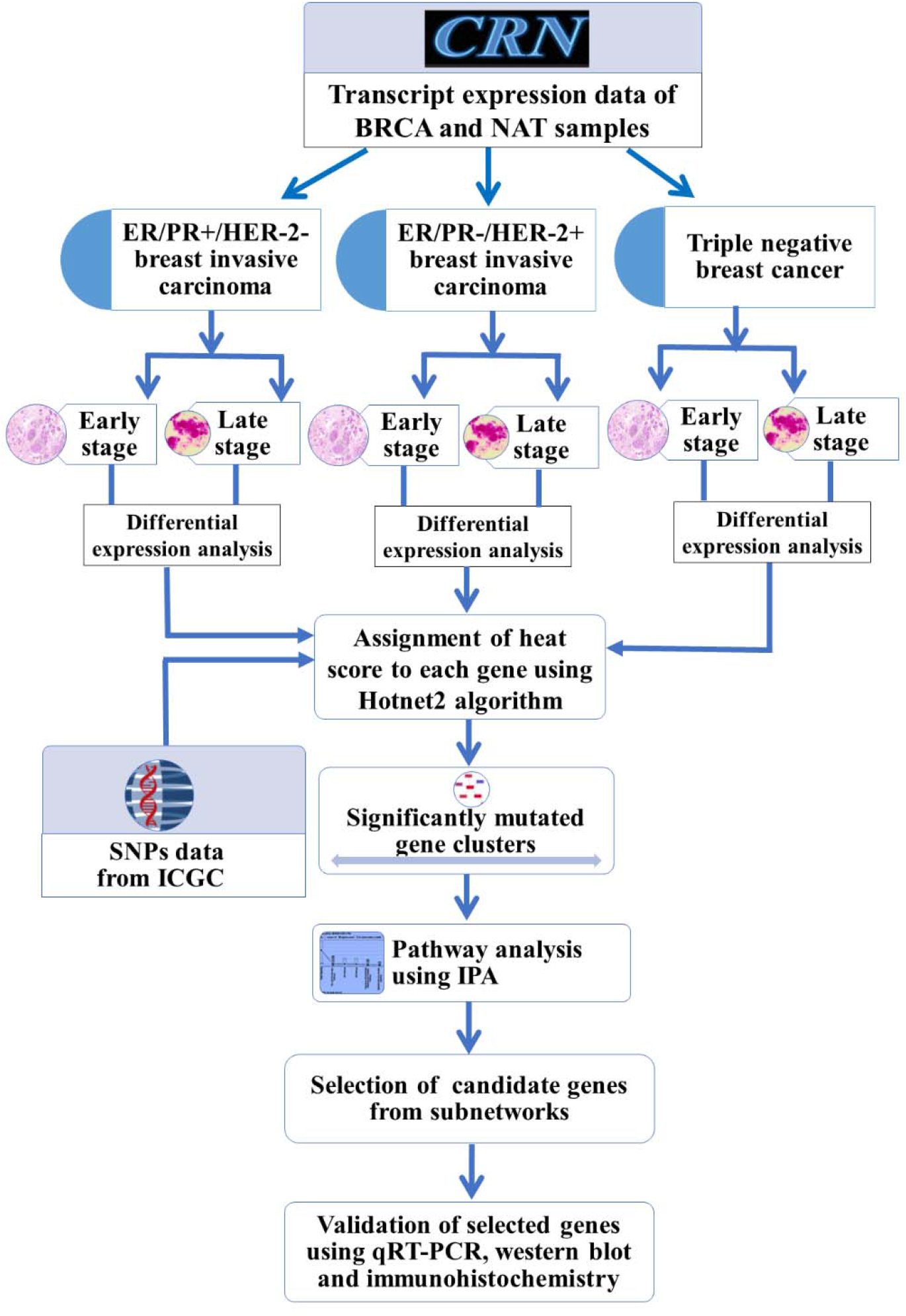
The flowchart of methodology. The Cancer RNA-seq Nexus (CRN) and International Cancer Genome Consortium (ICGC) were used as data sources as indicated in the figure. The differential expression analysis was performed on BRCA (breast invasive carcinoma) and NAT (Normal Adjacent Tissue) samples in three major molecular classes of BRCA in early and late stage. The genes were clustered into subnetworks based on their frequency of mutation. The pathway and functional enrichment analysis were performed on selected clusters of genes and selected genes were validated using qRT-PCR, western blot and immunohistochemistry. IPA=Ingenuity Pathway Analysis. qRT-PCR=Quantitative Real-Time Polymerase Chain Reaction. SNP=Single Nucleotide Polymorphism.

**Figure 2.**
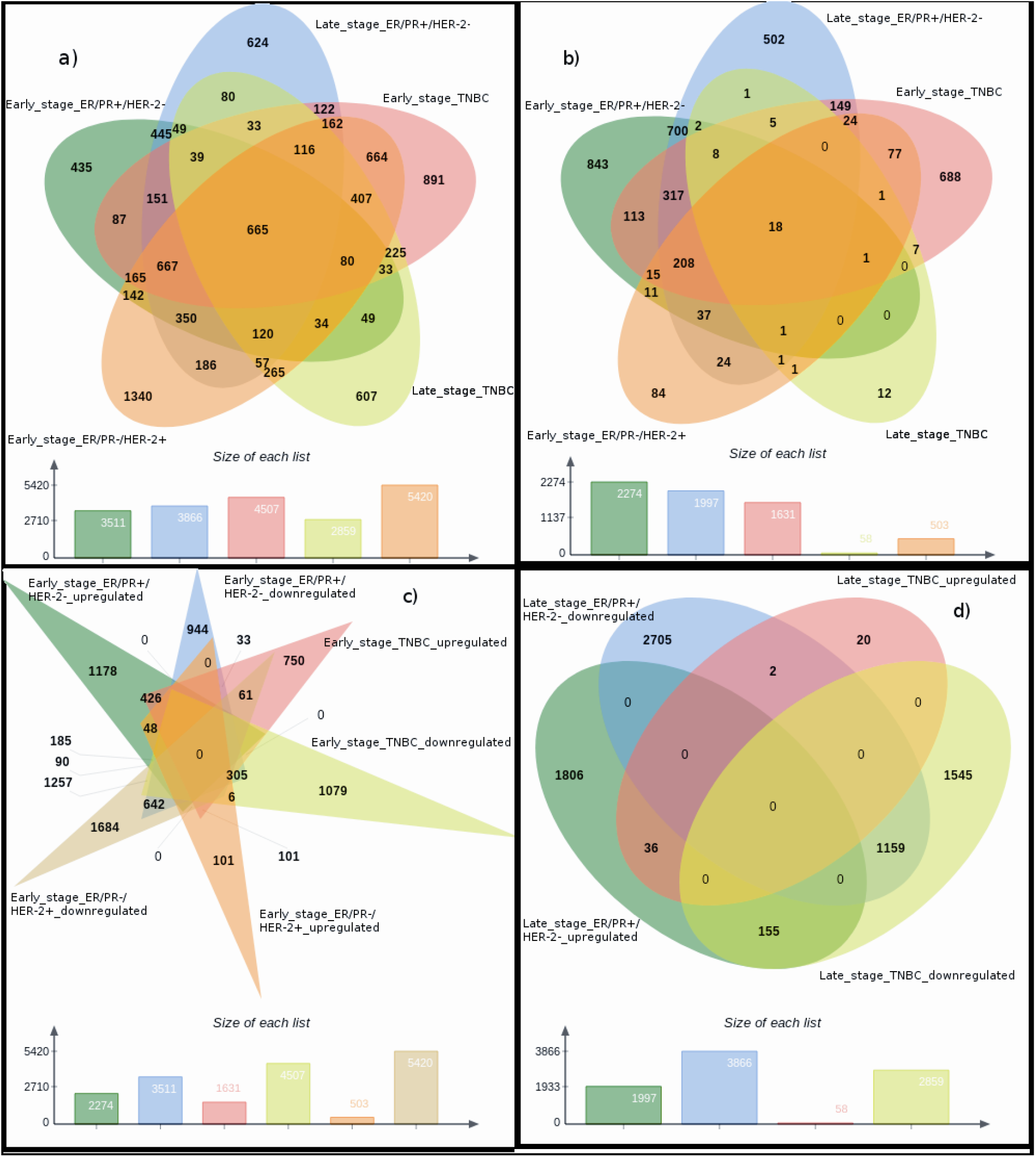
The dysregulated genes among different classes a) Stage specific downregulated genes among different classes. Total 665 genes were found to be downregulated in all categories i.e. Early stage ER/PR+/HER-2-; Late stage ER/PR+/HER-2-; Early stage ER/PR-/HER-2+; Early stage TNBC and Late stage TNBC. b) the overlap of upregulated genes among different classes. The 18 genes were found to be commonly upregulated in all five classes Early stage ER/PR+/HER-2-; Late stage ER/PR+/HER-2-; Early stage ER/PR-/HER-2+; Early stage TNBC and Late stage TNBC. Commonly dysregulated genes in early (c) and late stage (d) The inverse expression pattern was accorded for 38 genes which were downregulated in Early stage ER/PR+/HER-2- and upregulated in early stage TNBC. Similarly, genes showed inverse patterns among other classes as well. The detailed information is given in table 3 and supplementary table S2a, S2b and S2c.

**Figure 3.**
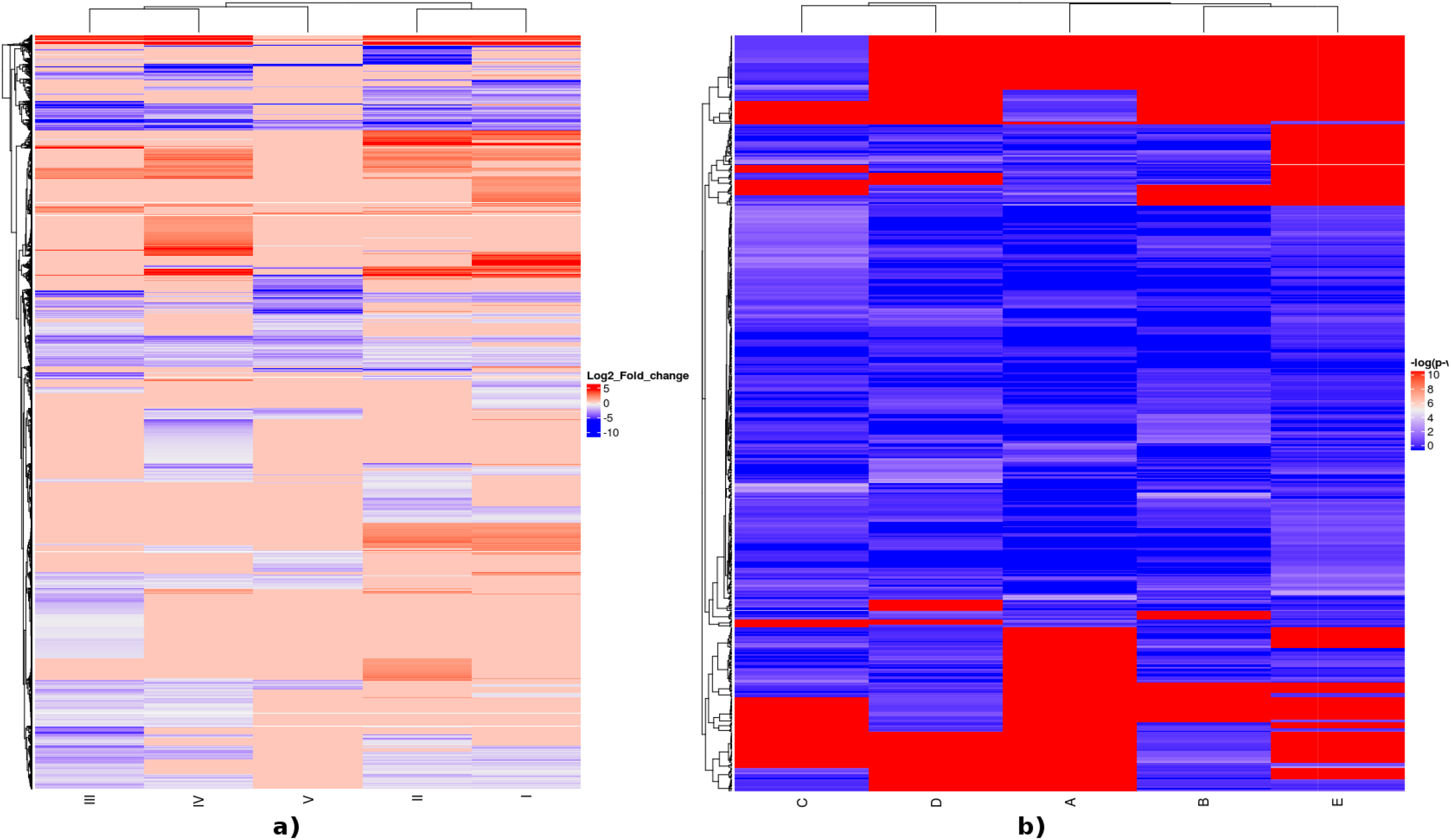
a) Heatmap showing significant biological pathways in five categories i.e. early stage ER/PR+/HER-2-, late stage ER/PR-/HER-2+, early stage ER/PR-/HER-2+, early stage TNBC and late stage TNBC. The colour scale represents -log(p-value) from 0 to 10. b). A= early stage ER/PR+/HER-2-, B= early stage ER/PR-/HER-2+, C= early stage TNBC, D=late stage ER/PR-/HER-2+, and E=late stage TNBC. Heatmap representing differentially expressed genes among all classes. The color scale represents log_2_ fold change values. The blue color represents downregulation of genes while red color represents upregulation of genes. I= early stage ER/PR+/HER-2-, II=late stage ER/PR-/HER-2+, III=early stage ER/PR-/HER-2+, IV=early stage TNBC and V=late stage TNBC.

**Table 1.**
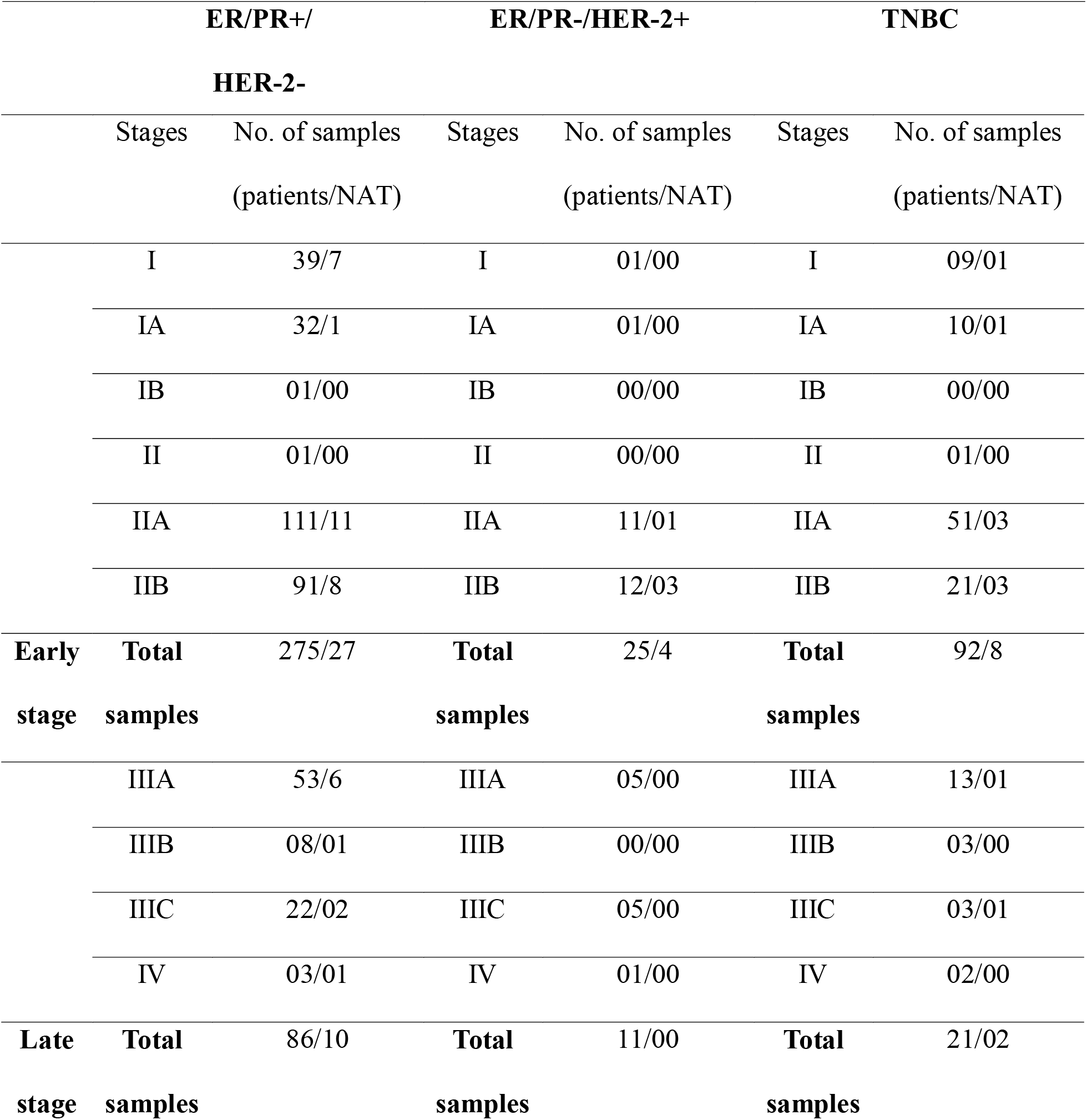
Summary of the number of samples obtained for BRCA (breast invasive carcinoma) and NAT (normal adjacent tissue) in each stage of ER/PR+/HER-2-, ER/PR- /HER-2+ and TNBC (triple negative breast cancer).

**Table 2.**
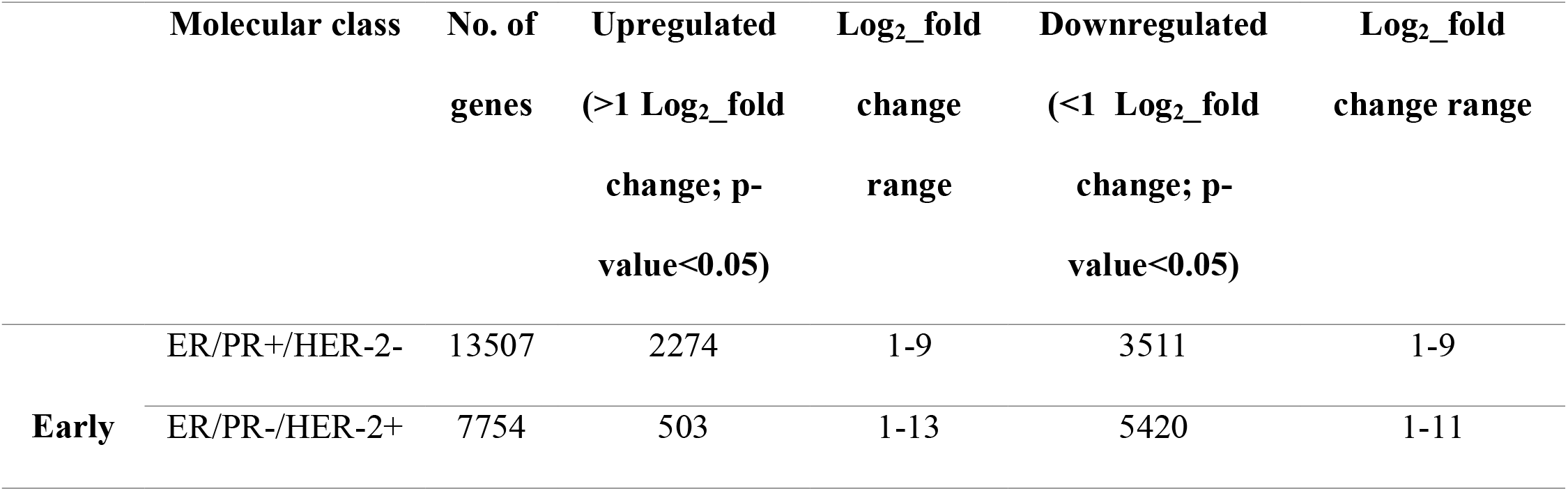
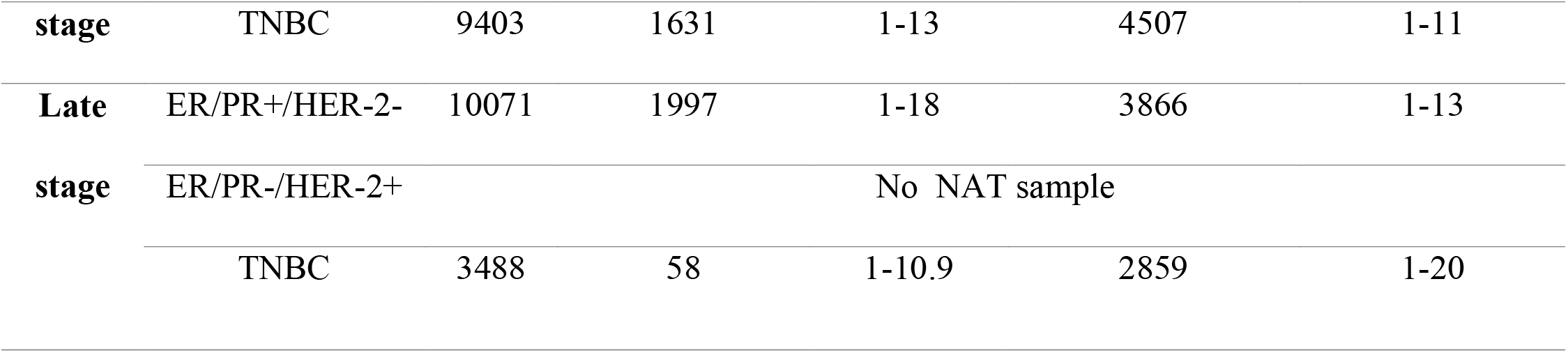
Summary of dysregulated genes in early and late stage in each of the three molecular classes. The table shows significantly dysregulated genes with a log_2_ fold change greater than or less than 1. In early stage, there was a greater number of dysregulated genes as compared to late stage. The log_2_ fold change varied from 1-13 in early stages and 1-20 in late stages. In the case of late-stage ER/PR-/HER-2+, the analysis could not be done as there was no sample of NAT to compare with. The larger number of genes were found to be downregulated in all classes.

|  | **Molecular class** | **No. of genes** | **Upregulated (>1 Log<sub>2</sub>_fold change; p-value<0.05)** | **Log<sub>2</sub>_fold change range** | **Downregulated (<1 Log<sub>2</sub>_fold change; p-value<0.05)** | **Log<sub>2</sub>_fold change range** |
| --- | --- | --- | --- | --- | --- | --- |
| **Early** | ER/PR+/HER-2- | 13507 | 2274 | 1-9 | 3511 | 1-9 |
|  | ER/PR-/HER-2+ | 7754 | 503 | 1-13 | 5420 | 1-11 |
| **found to be downregulated in all classes.** |  |  |  |  |  |  |
| **stage** | TNBC | 9403 | 1631 | 1-13 | 4507 | 1-11 |
| **Late** | ER/PR+/HER-2- | 10071 | 1997 | 1-18 | 3866 | 1-13 |
| **stage** | ER/PR-/HER-2+ |  |  | No NAT sample |  |  |
|  | TNBC | 3488 | 58 | 1-10.9 | 2859 | 1-20 |

**Table 3.**
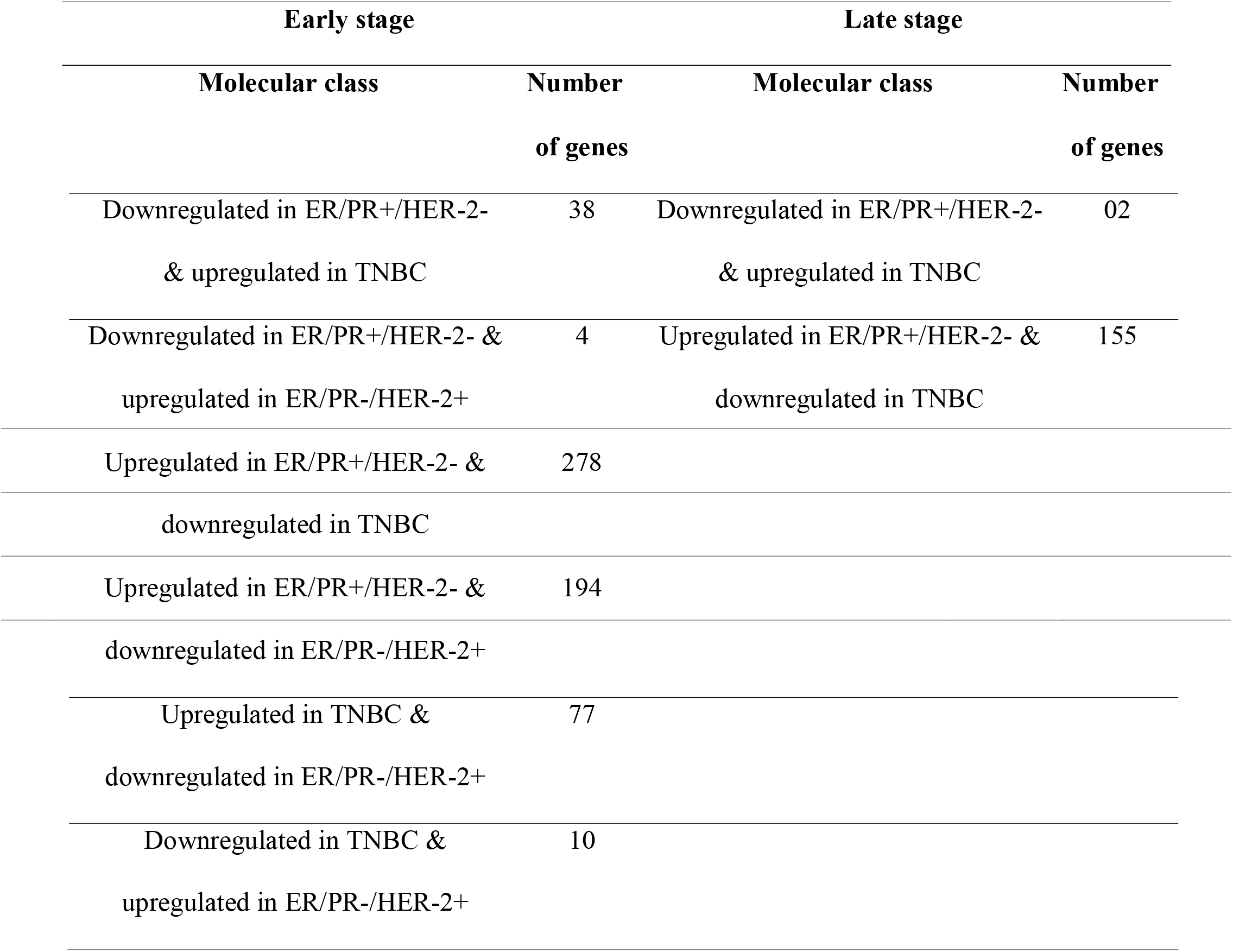
The genes showed inverse expression pattern among different classes in early and late stage. The genes which showed upregulation in one class were found to be downregulated in other class and vice-versa. The table indicates the number of such genes.

### Mutated subnetworks of genes

The differentially expressed genes were clustered into subnetworks based on PPIs data and number of single nucleotide polymorphisms (SNPs) they harbored, using hotnet2 algorithm[5]. A total of nine significantly mutated clusters were found in different molecular classes (table 4) containing hot (frequently mutated) and cold (less mutated) genes. The cold genes are implicated because of their interaction with hot genes (guilt-by-association). Pathway analysis indicated that some of them have role in well-known cancer signaling pathways while others are less characterized.

**Table 4.**
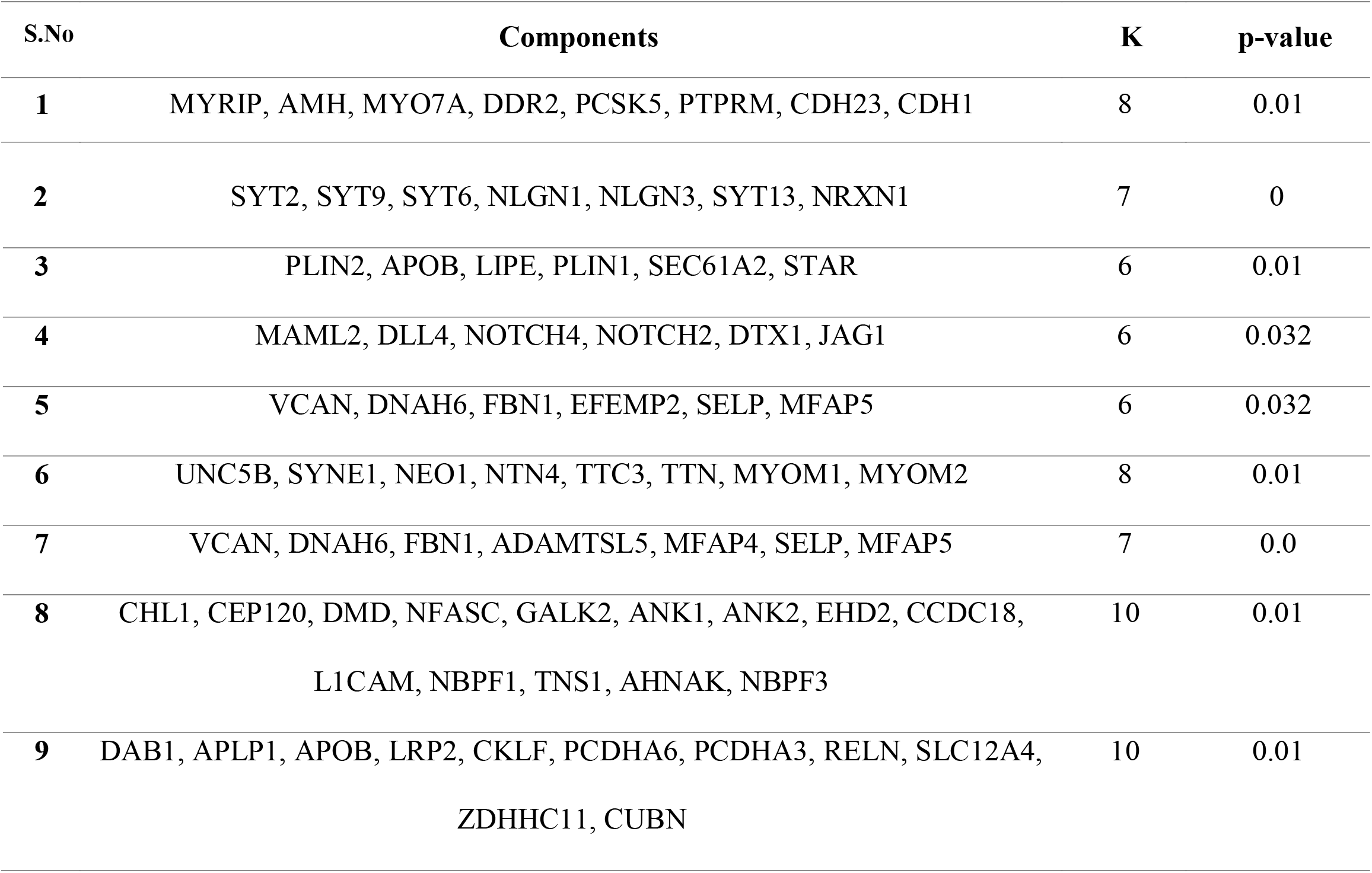
Summary of results obtained after hotnet2 analysis. For each subnetwork, the
table shows the minimum K for which subnetworks of size ≥ K are significant (p-value < 0.05).

For each of the categories, different clusters were obtained that are discussed class-wise.

### Early stage ER/PR+/HER-2-

Three mutated clusters were obtained, out of which one is significantly mutated (p-value<0.05). containing seven genes viz. SYT2, SYT6, SYT9, SYT13, NRXN1, NLGN1 and NLGL3 (Fig. 4a). Among them, 2 were upregulated and 5 were downregulated with average log_2_ fold change (FC) as 2.21. The details are given in table 5a (FC, heat score, pathways etc.). Their differential expression was further examined in online databases viz. Oncomine [6] and Expression atlas [7] in BRCA. All of them were found to be differentially expressed except NLGN1 (Table 5a, supplementary table S4). These results are largely in-line with our observations. The pathway analysis using IPA indicated that the cluster may be involved in TR/RXR activation pathway (supplementary fig. S1), which activates PI3K pathway and is involved in cell survival and growth [8]. The associated GO terms (biological processes, molecular function and cellular component) are given in supplementary table S5a.

**Figure 4.**
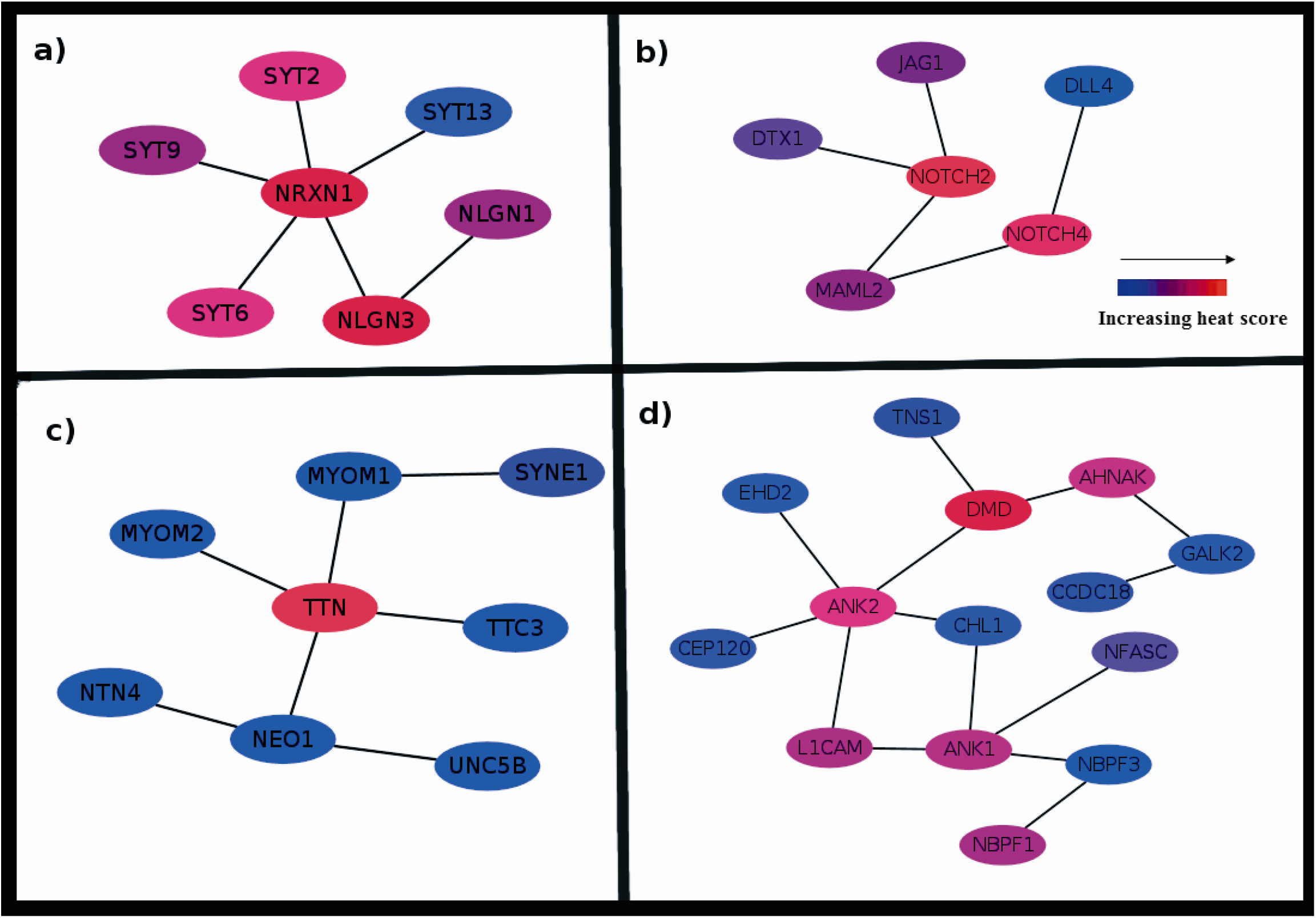
Subnetwork for (a) Early stage ER/PR+/HER-2-. Seven dysregulated genes, coloured by the increasing mutation frequency (blue to red). NLGN3 and NRXN1 are frequently mutated while SYT13 is less mutated. The pathway analysis showed enrichment of TR/RXR activation pathway which activates PI3K pathway and regulates the biological process involved in cell survival and cell growth. (b) Early stage ER/PR-/HER-2+ containing six dysregulated genes. The NOTCH2 and NOTCH4 are highly mutated. The subnetwork may be involved in Notch signaling pathway, Th1 and Th2 activation pathway and epithelial-mesenchymal transition pathway. (c) Early stage TNBC. Contains eight dysregulated genes among which TTN is highly mutated which is followed by SYNE1. Netrin signaling, RhoA signaling, and axonal guidance signaling pathways may be affected. (d) Late stage TNBC. Contains fourteen dysregulated genes. DMD2 and ANK2 are significantly enriched in mutations. The subnetwork may regulate galactose degradation I pathway, colanic acid building blocks biosynthesis and nNOS signaling pathway.

**Table 5a.**
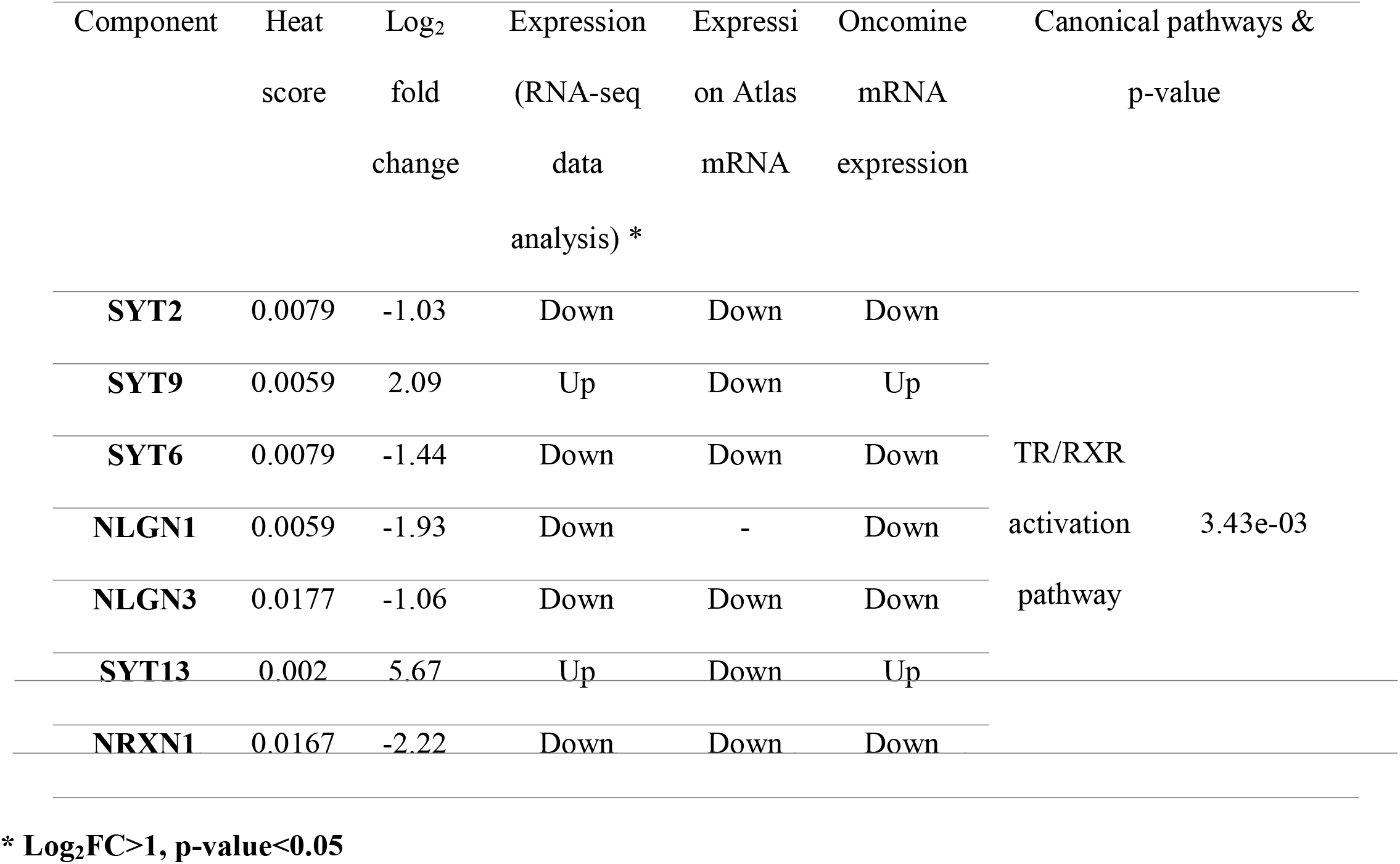
Summary of each component of the subnetwork. The table is showing heat score and fold change of each component. The status and mRNA expression of the components in expression atlas and Oncomine is also shown.

### Early stage ER/PR-/HER-2+

Two significant clusters were identified and most significant is discussed which contained 6 downregulated genes viz. JAG1, NOTCH2, DTX1, MAML2, NOTCH4 and DLL4 (average log_2_ FC=1.60, Fig. 4b, table 5b). A search in Oncomine and Expression Atlas revealed all of them to be downregulated (Table 5b, supplementary table S4). The cluster was found to be involved in Notch signaling, Th1 and Th2 activation and regulation of epithelial-mesenchymal transition (EMT) pathways (supplementary fig. S2a, S2b, S2c). which are reported to be involved in causation and progression of BRCA. The NOTCH and its receptors are involved in cell proliferation, differentiation, survival and progression of BRCA [9–11]. Inflammatory biomarkers (Th1, Th2) may be linked to risk of BRCA carcinogenesis [12]. Epithelial to mesenchymal transition (EMT) facilitates carcinoma cells to gain mobility and they migrate from primary to other sites, which can lead to invasion and metastasis in cancer cells [13]. The GO terms associated with this cluster are given in supplementary table S5b.

**Table 5b.**
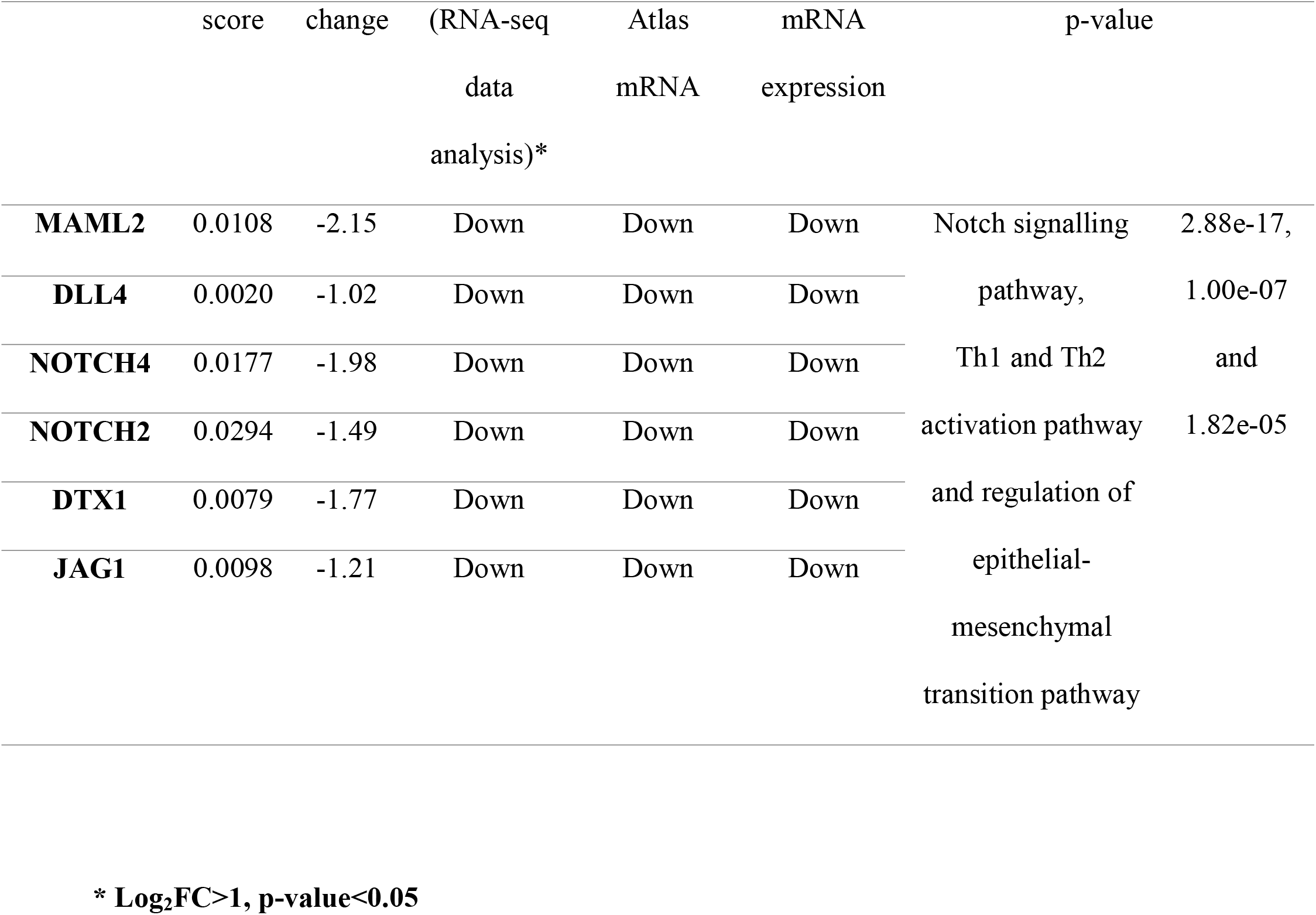
Summary of each component of the subnetwork. The heat score value of each component is calculated based on the mutation frequency. The subnetwork is may be involved in pathways e.g. Notch signaling pathway, Th1 and Th2 activation pathway and regulation of epithelial-mesenchymal transition pathway (EMT).

### Early stage TNBC

Two mutated subnetworks were identified and most mutated composed of 8 genes viz. TTN, TTC3, MYOM1, MYOM2, SYNE1, NEO1, NTN4 and UNC5B (average log_2_ FC=2.17). All genes in subnetwork were significantly downregulated except UNC5B which was upregulated. In Oncomine, UNC5B, NEO1 and TTC3 were found to be upregulated and remaining were found to be downregulated. The expression atlas showed UNC5B and TTN to be upregulated and rest to be downregulated (table 5c and supplementary table S4). The subnetwork contained both hot genes and cold genes (Fig. 4c). TTN and SYNE1 are known driver genes in BRCA and were significantly mutated as compared to others and [14]. Pathway analysis showed enrichment of netrin signaling, RhoA signaling, and axonal guidance signaling pathways (supplementary fig. S3a, S3b, S3c). Netrin-1 is overexpressed in large fraction of MBC (metastatic breast cancer) [15] and is a potential therapeutic target. Inhibition of RhoA signaling pathway inhibits breast cancer cell invasion [16]. Axonal guidance signaling molecules play a crucial mechanism in tumor suppression and oncogenesis and can be promising targets for therapeutic development [17, 18].

**Table 5c.** Summary of each component of the subnetwork in early stage TNBC class. The table also depicts status and mRNA expression of each component in expression atlas and Oncomine.

| Component | Heat<br>score | Fold<br>change | Expression<br>(RNA-seq<br>data<br>analysis) * | Expression<br>Atlas<br>mRNA | Oncomine<br>mRNA<br>expression | Canonical pathways &<br>p-value |  |
| --- | --- | --- | --- | --- | --- | --- | --- |
| **UNC5B** | 0.0088 | 1.58 | Up | Up | Up | Netrin | 2.61e-02, |
| **SYNE1** | 0.0558 | -1.68 | Down | Down | Down | signalling, | 4.85e-02 |
| **NEO1** | 0.0088 | -1.11 | Down | Down | Up | RhoA | and |
| **NTN4** | 0.0049 | -2.87 | Down | Down | Down | signalling, | 1.33e-02 |
| **TTC3** | 0.0127 | -1.17 | Down | Down | Up | and axonal |  |
| **TTN** | 0.2008 | -2.90 | Down | Up | Down | guidance |  |
| **MYOM1** | 0.0078 | -4.15 | Down | Down | Down | signaling |  |
| **MYOM2** | 0.0147 | -1.90 | Down | Down | Down | pathways |  |

**\* Log<sub>2</sub>FC>1, p-value<0.05**

### Late stage TNBC

Two mutated clusters were found and most significant is comprised of 14 genes viz. TNS1, DMD, AHNAK, GALK2, CCDC18, ANK2, EHD2, CHL1, CEP120, ANK1, NFASC, LICAM, NBPF3 and NBPF1 (Fig. 4d) (average log_2_ FC 2.59). It indicates majority of them are highly up or downregulated. A search in Oncomine database indicated that CCDC18, NBPF3, NFASC, TNS1 and L1CAM was upregulated while the rest were found downregulated. In the expression atlas CCDC18, NBPF3, NBPF1 and ANK1 were found to be upregulated whereas rest were downregulated (Table 5d and supplementary table S4). It is important to note that TNS1, DMD and ANK2 [14] and AHNAK [19] are known driver genes in BRCA. The cluster was found to be involved in galactose degradation I, colanic acid building blocks biosynthesis and nNOS signaling pathway (supplementary fig. S4a, S4b, S4c). It has been reported that knockdown of galactokinase gene control growth of hepatoma cells in vitro [20] and galactose metabolism pathways are dysregulated in breast cancer [21]. However, there was no literature report of colanic acid building blocks biosynthesis and nNOS signaling pathway in BRCA. The associated GO terms are given in supplementary table S5c.

**Table 5d.**
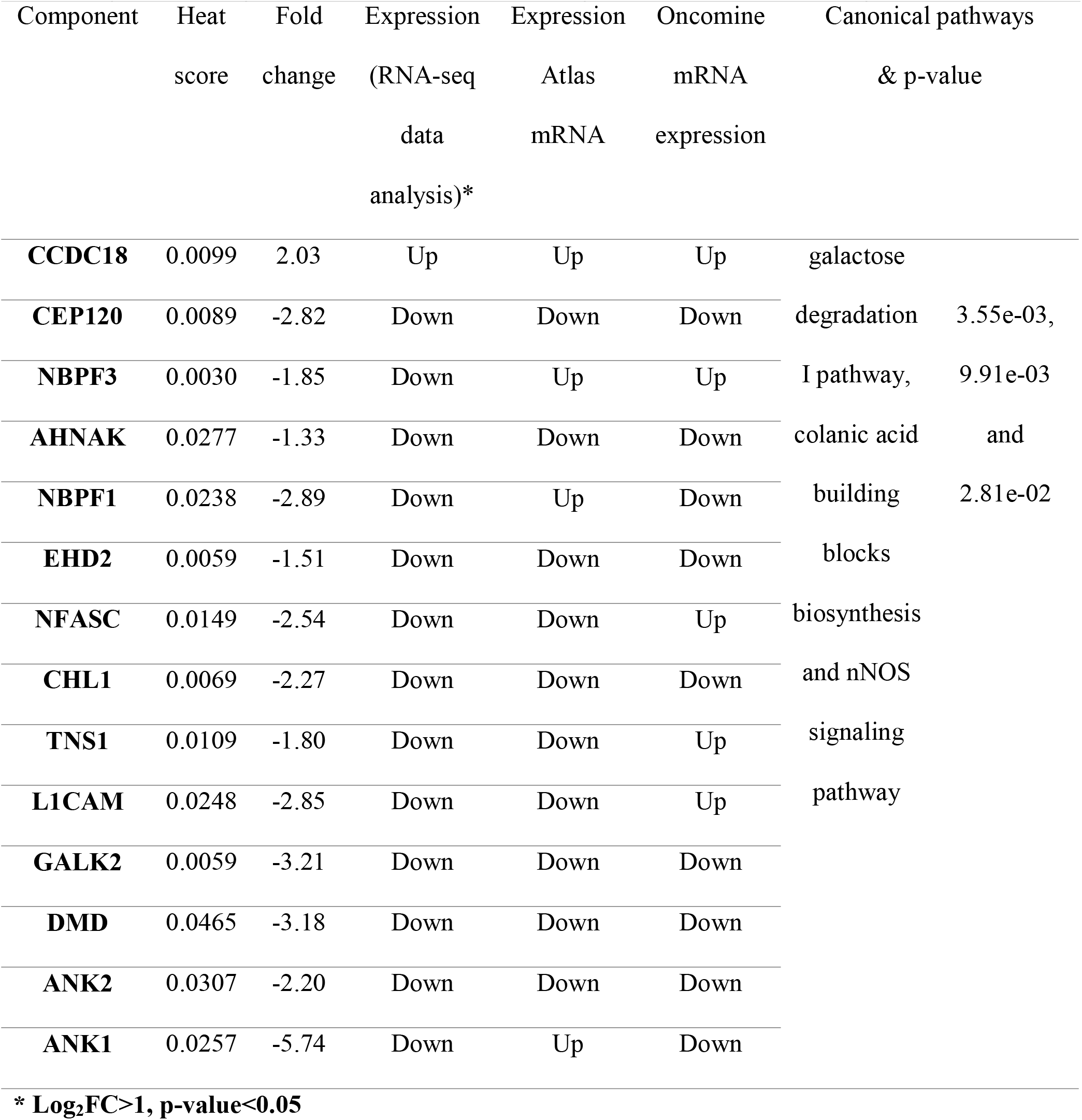
Summary of each component of the subnetwork in late stage TNBC. The table also represents status and mRNA expression of each component in expression atlas and Oncomine. The subnetwork appears to be involved in pathways as galactose degradation I, colanic acid building blocks biosynthesis and nNOS signaling pathway.

The cluster from all classes contain 35 genes in total containing less mutated genes associated with frequently mutated genes. Many of these gene (e.g. NOTCH2, NOTCH4, DMD [22], CCDC18 [23], EHD2 [24], JAG1 [25] are known to play important roles in BRCA and other cancers.

It was interesting to see many genes in the identified clusters which are not yet reported to be directly involved in BRCA. An initial analysis indicated that these genes have reported interactions with known cancer genes. Some of them are regulators of genes known for their role in BRCA. Their GO terms, pathways, biological functions, binding and regulations have been discussed later. Additionally, many of them were also found to be dysregulated across other cancers [26] (Fig. 5). Therefore, it is reasonable to assume that they may also play some role in BRCA. The genes were further examined for inframe, truncating, missense or other mutations using cBioPortal [27]. The data was obtained from four studies for breast invasive carcinoma (TCGA, Cell 2015, TCGA, Nature 2012, TCGA, PanCancer Atlas and TCGA, Provisional). A large number of mutations were found in different genes viz. TTN, SYNE1, ANK2, and NLGN3. In fact, few genes showed mutation frequency greater than BRCA1 and BRCA2.

**Figure 5.**
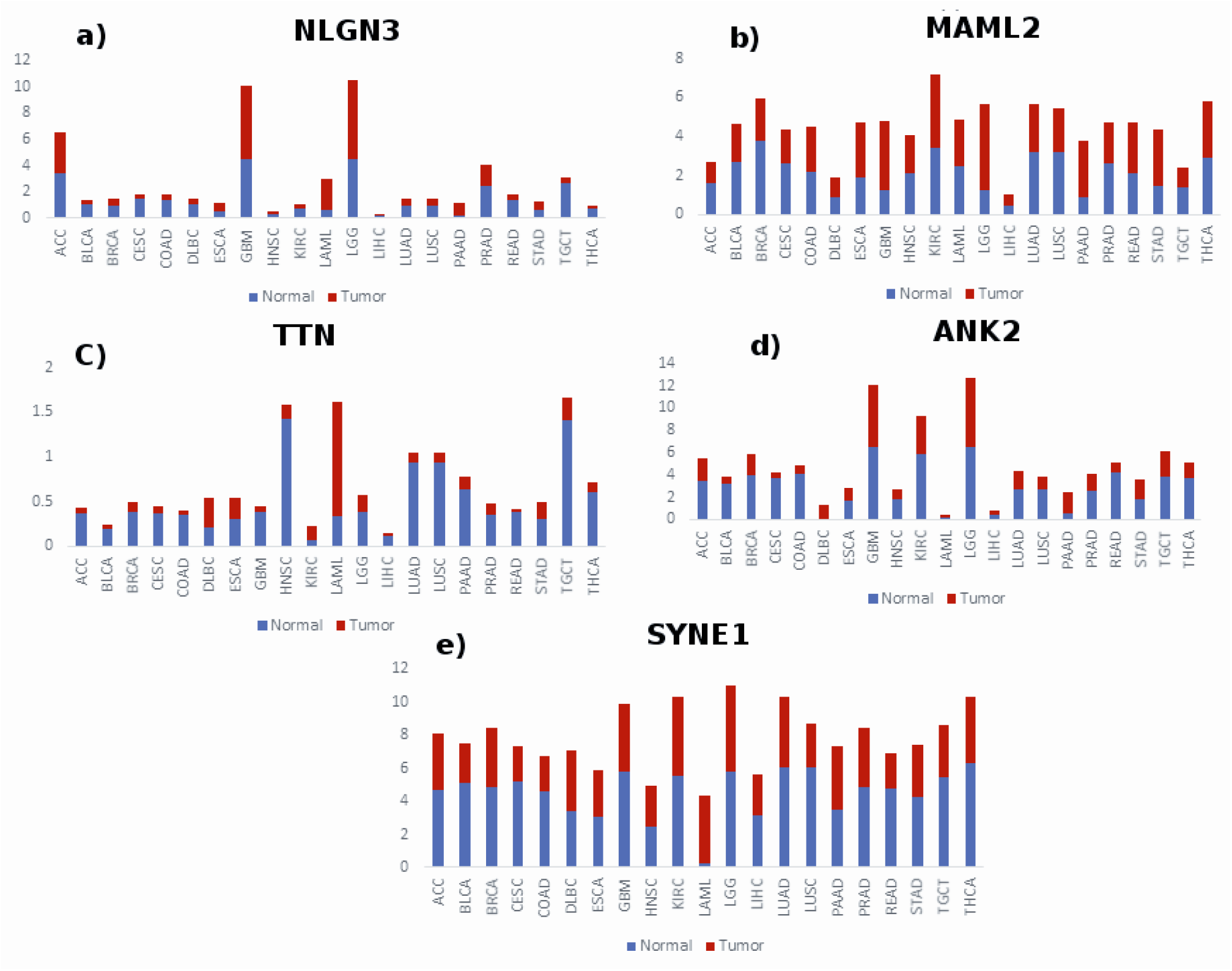
Expression profile of selected genes in the tumor (red) and normal tissues (blue) across different cancers. The X-axis represents normal and tumor samples and Y-axis represents median expression (p-value ≤ 0.01). The median expression values are the log_2_(TPM + 1) values. ACC; Adrenocortical carcinoma, BLCA; Bladder Urothelial Carcinoma, BRCA; breast invasive carcinoma, CESC; cervical squamous cell carcinoma and endocervical adenocarcinoma, COAD; colon adenocarcinoma, DLBC; Lymphoid Neoplasm Diffuse Large B-cell Lymphoma, ESCA; esophageal carcinoma, GBM; Glioblastoma multiforme, HNSC; Head and neck squamous cell carcinoma, KIRC; Kidney renal clear cell carcinoma, LAML; Acute Myeloid Leukemia, LGG; Brain Lower Grade Glioma, LIHC; liver hepatocellular carcinoma, LUAD; lung adenocarcinoma, LUSC; lung squamous cell carcinoma, PAAD; Pancreatic adenocarcinoma; PRAD; Prostate adenocarcinoma, READ; Rectum adenocarcinoma, STAD; stomach adenocarcinoma, THCA; Thyroid carcinoma, TGCT; Testicular Germ Cell Tumors, TPM; Transcripts per Million.

### Mutational analysis of clusters

#### Early stage ER/PR+/HER-2-: NLGN3, NRNX1, NLGN1 cluster

In NLGN1 and NLGN3, carboxylesterase domain contains majority of the mutations whereas the laminin G domain contains majority of mutations in NRXN1. The mutations laminin domain may contribute to EMT via affecting cell-to-cell adhesion with binding to integrin α6ß4 (ß4 subunit) expressed by epithelial cells. This promote tumor cell survival, invasion and migration by signaling through RAC and PI3K pathway [28]. In other members (SYT2, SYT6, SYT9 and SYT13), C2 domain was affected. The C2 domains of SYT family are involved in exocytosis. The alterations of exocytic proteins are known to be associated with malignant transformations [29]. The altered exocytic networks are involved in the acquisition of prometastatic traits [30]. Therefore, the identification of such a cluster is no surprise. The details of the mutations in each gene are given in table 6a and supplementary table S6a. Such mutations which can impact protein function have been plotted against tumour samples (Fig. 6a). The deleterious mutations were identified using PolyPhen-2, Mutation Assessor and SIFT algorithm. The percent mutation rate for each gene was computed using number of deleterious mutations it carries. Among all genes in this subnetwork, the NLGN3 showed higher mutation rate, equal to BRCA1.

**Figure 6.**
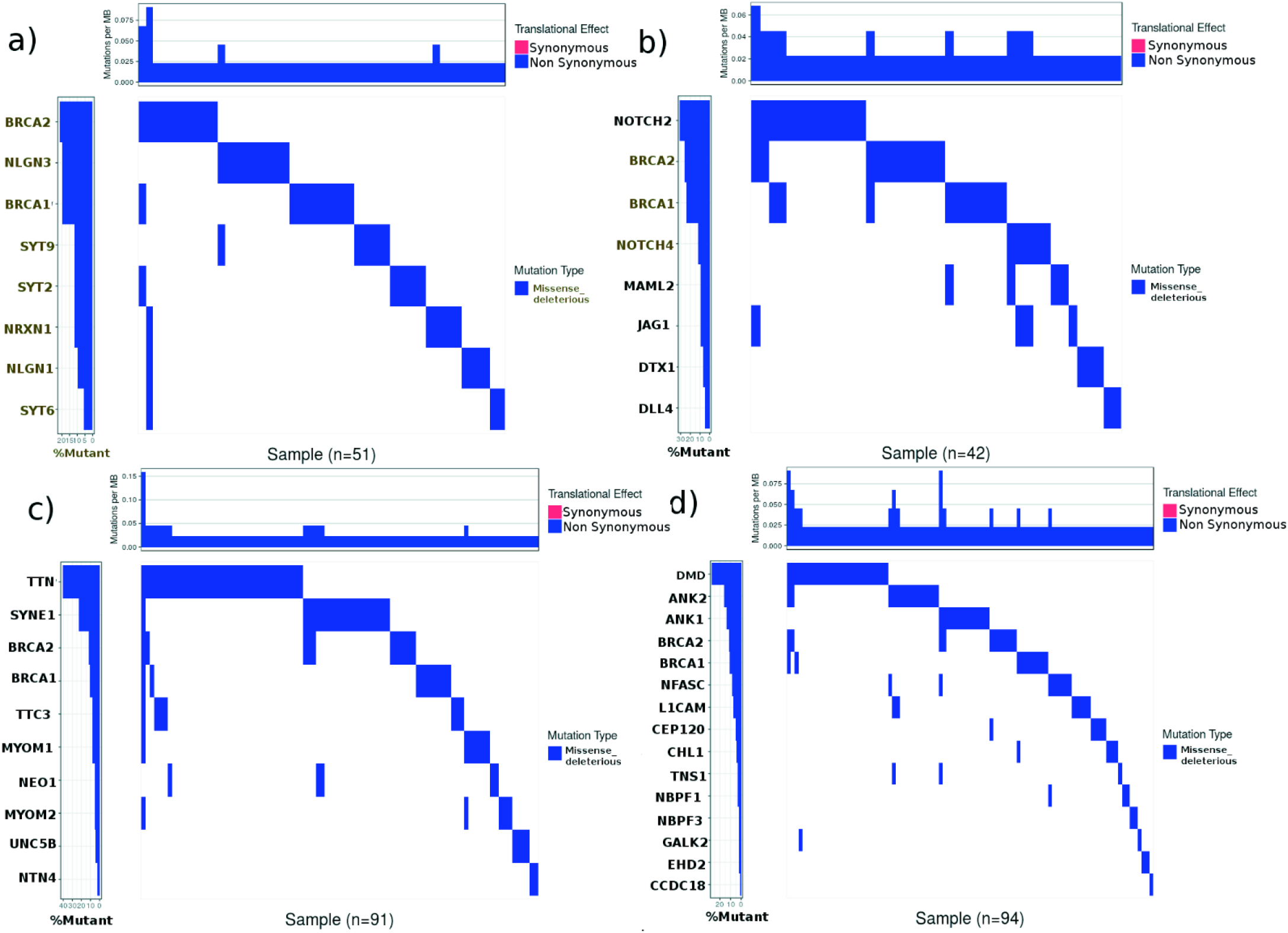
Mutational landscape of candidate genes showing deleterious functional impact in (a) 51 out of 60 samples, (b) 42 out of 59 samples (c) 91 out of 254 samples (d) 94 out of 164 samples. For each sample, the mutations per MB was calculated using number of deleterious mutations present. The mutation rate for a gene is based on deleterious missense mutation in number of samples. The mutation rate in NLGN3 is equal to BRCA1 (10 deleterious mutations). The mutation rate in TTN, SYNE1, ANK1 and ANK2 is greater than BRCA1 and BRCA2.

**Table 6a.**
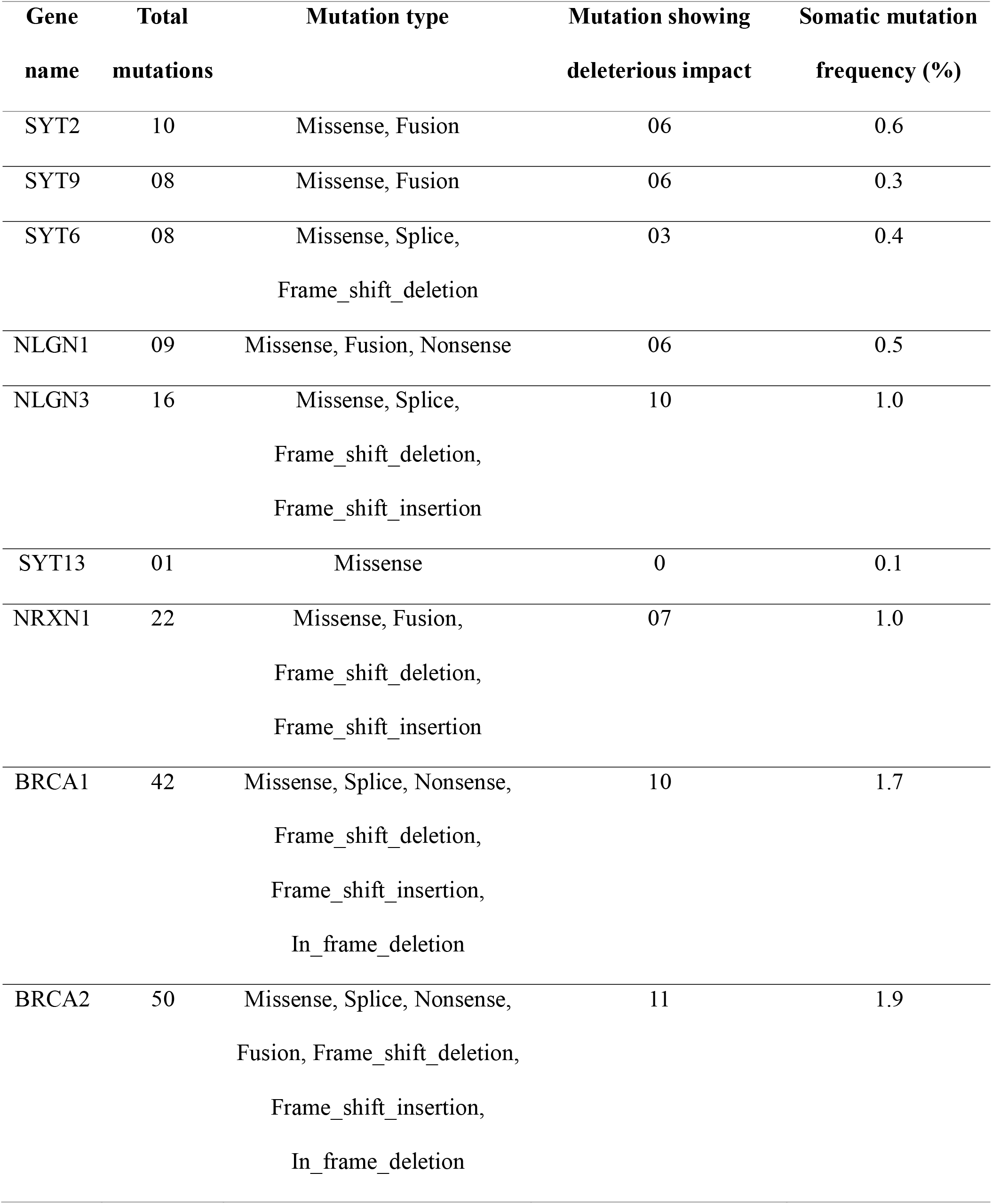
Mutational detail of the components of subnetwork showing total mutations (unique) across the four studies, TCGA; Cell 2015, TCGA; Nature 2012, TCGA; PanCancer Atlas and TCGA; Provisional using cbioportal. The cluster harbored several types of mutations as missense, fusion, splice, frame-shift, in-frame. The somatic mutation frequency of NLGN3 and NRXN1 was found to 1.0%.

**\* Log<sub>2</sub>FC>1, p-value<0.05**
| Gene | Total | Mutation type | Mutation showing | Somatic mutation |
| --- | --- | --- | --- | --- |
| name | mutations |  | deleterious impact | frequency (%) |
| SYT2 | 10 | Missense, Fusion | 06 | 0.6 |
| SYT9 | 08 | Missense, Fusion | 06 | 0.3 |
| SYT6 | 08 | Missense, Splice,<br>Frame_shift_deletion | 03 | 0.4 |
| NLGN1 | 09 | Missense, Fusion, Nonsense | 06 | 0.5 |
| NLGN3 | 16 | Missense, Splice,<br>Frame_shift_deletion,<br>Frame_shift_insertion | 10 | 1.0 |
| SYT13 | 01 | Missense | 0 | 0.1 |
| NRXN1 | 22 | Missense, Fusion,<br>Frame_shift_deletion,<br>Frame_shift_insertion | 07 | 1.0 |
| BRCA1 | 42 | Missense, Splice, Nonsense,<br>Frame_shift_deletion,<br>Frame_shift_insertion,<br>In_frame_deletion | 10 | 1.7 |
| BRCA2 | 50 | Missense, Splice, Nonsense,<br>Fusion, Frame_shift_deletion,<br>Frame_shift_insertion,<br>In_frame_deletion | 11 | 1.9 |

#### NLGN3 as a candidate gene in early stage ER/PR+/HER-2-

The fact that the NLGN3 is dysregulated and highly mutated in number of samples makes it interesting candidate. NLGN3 is protein-coding gene, that is reported to be involved in Asperger’s syndrome and autism X-linked 1. The gene was found to be significantly downregulated (log: FC −1.06) (supplementary fig. S5a) and interacts with other genes viz. DLG4, GABRA1, OR1F12, CYFIP2, GABRA2, AGAP2, NLGN4X, UTP15, NLGN4Y, TMEM39A, TYK2, RECQL4, PIK3CB, PIK3CA, NLGN1 [31]. Many of them are well-known cancer associated genes. The NLGN3 also found to have a role in cell-cell adhesion and cell differentiation [32]. Additionally, it regulates GRIN2B [33], which is reported to play a significant role in cancer development and progression. In NLGN3, total sixteen mutations were found in different domains; among them ten were predicted deleterious. The figure 7a shows that carboxylesterase family, neurolgin-3 and alpha-beta hydrolase domain got significantly enriched in mutations. Therefore, considering the above reasons, the gene NLGN3 can be a good candidate gene.

**Figure 7.**
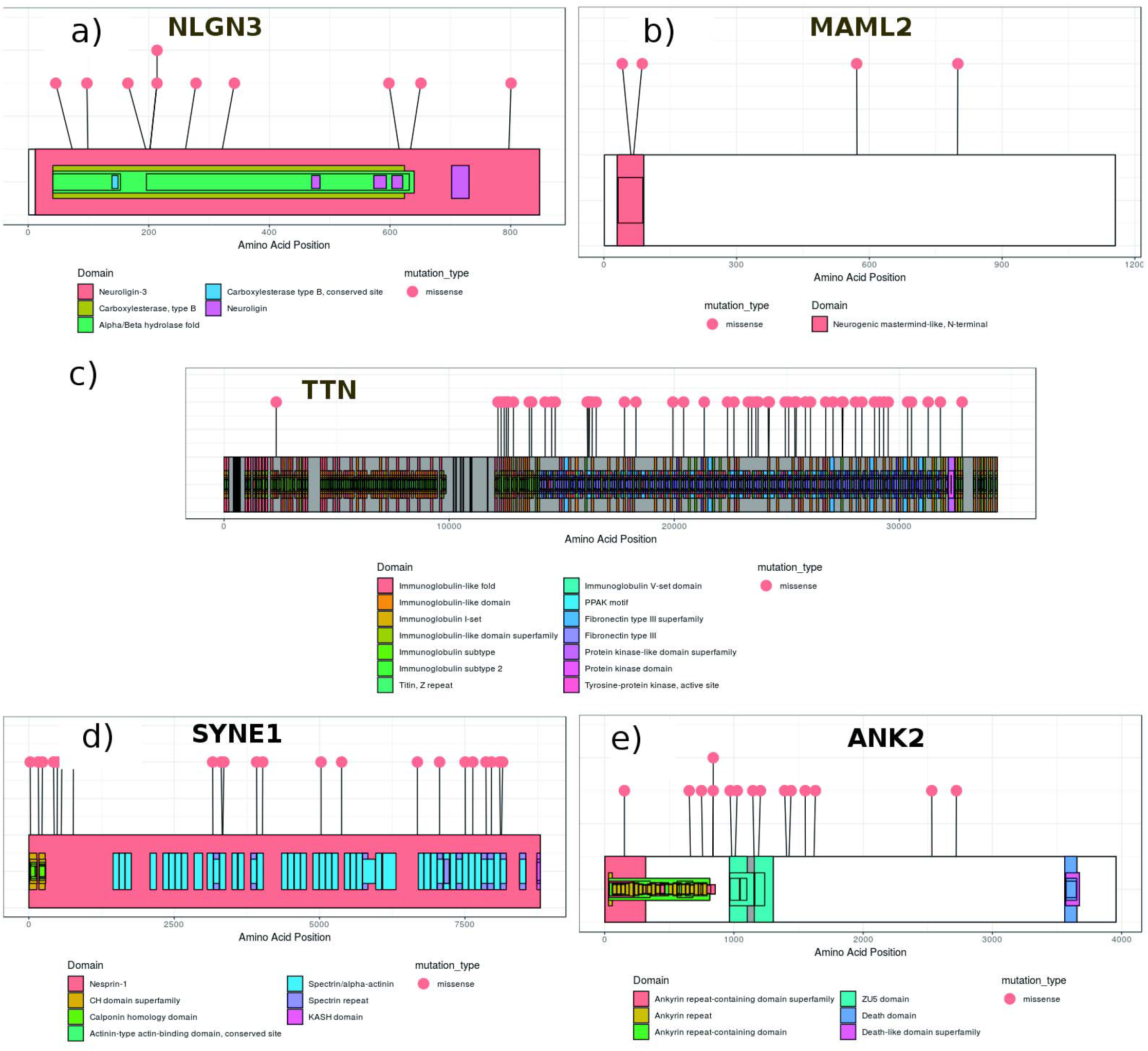
Lollipop plot representing missense mutations (deleterious) in functional domains of (a) NLGN3, (b) MAML2, (c) TTN, (d) SYNE1 and (e) ANK2. In NLGN3, carboxylesterase family, neurolgin-3 and alpha-beta hydrolase domain were significantly enriched in mutation. In MAML2, neurogenic mastermind like N-terminal domain harbored mutations as G61D and D67N. All functional domains in TTN were occupied with mutations. In SYNE1, the spectrin/alpha-actinin and spectrin-repeat domain got enriched and in ANK2, ZU5 domain got enriched.

#### Early stage ER/PR-/HER-2+: NOTCH and interactors

NOTCH2 and NOTCH4 are well-known genes in BRCA and other cancers. Most of the mutations in NOTCH2 and NOTCH4 were found to be missense that are spread over different domains viz. EGF-like, calcium binding EGF and NOTCH protein. Few truncating mutations also occurred in EGF-like domain and domain of unknown function (DUF). The mutations in EGF-like domain lead to inactivation of NOTCH [34]. The other members were less mutated and clustered due to their association with NOTCH. The details of different mutations in each of the gene is given in table 6b and supplementary table S6b. MAML2 harbored total 12 mutations, out of which, 4 were predicted to be deleterious. The figure 6b shows percent mutation rate for each member. Although, percent mutation rate for MAML2 is less, it interacts with NOTCH1 and NOTCH2, its role is not yet known in BRCA.

**Table 6b.**
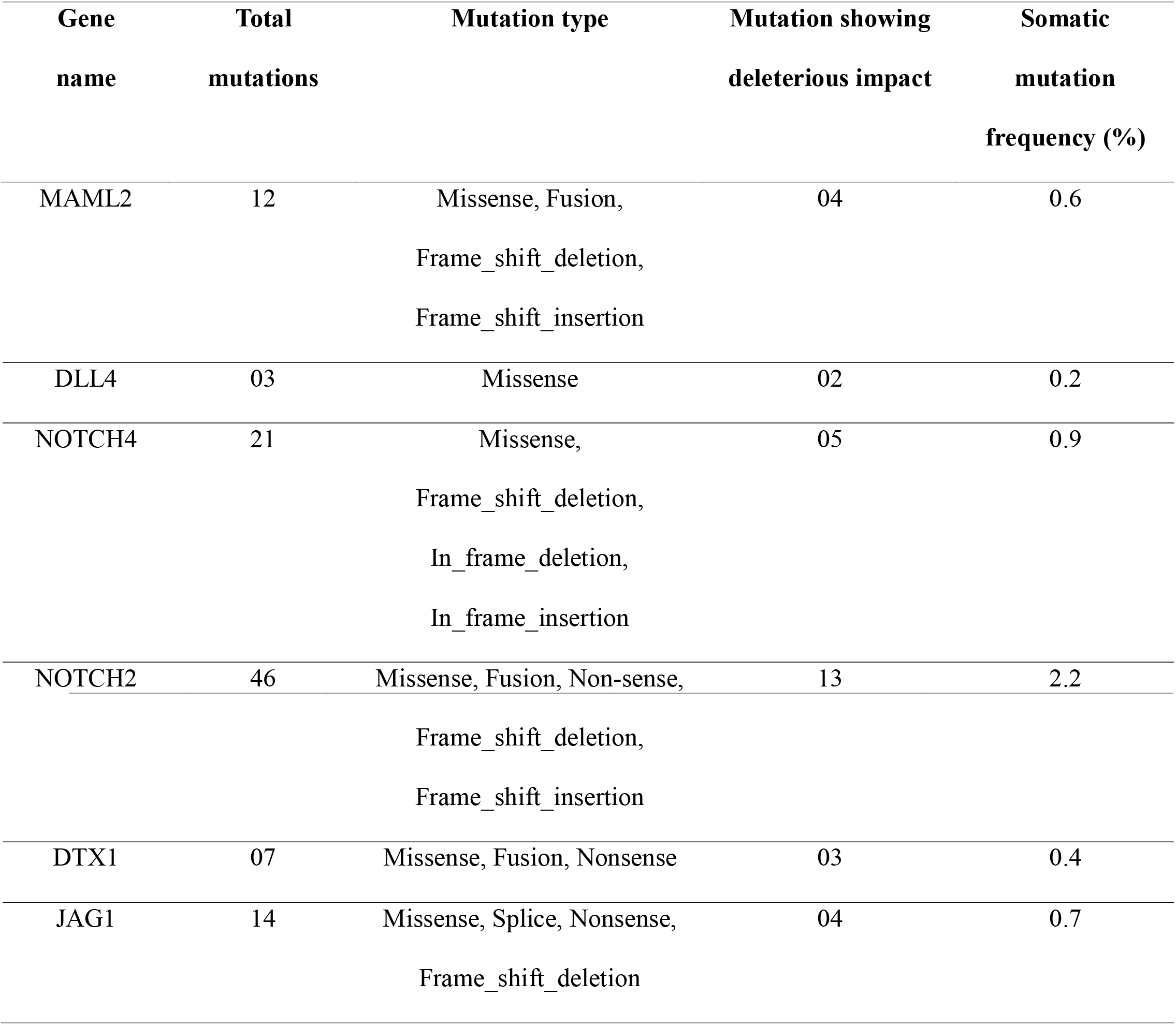
Mutational detail of the NOTCH gene and interactors representing total unique mutations across the four studies, TCGA; Cell 2015, TCGA; Nature 2012, TCGA; PanCancer Atlas and TCGA; Provisional. The components harbored several types of mutations as missense, fusion, splice, frame-shift, in-frame as shown in the table.

#### MAML2 as candidate gene in early stage ER/PR-/HER-2+

MAML2, a mastermind like transcriptional coactivator 2, encodes gene of mastermind-like family of proteins. It is known to be involved in hidradenoma of skin, adenolymphoma and carcinoma of the thymus. Its conserved domain present in N-terminus interacts with ankyrin repeat domain of NOTCH proteins (NOTCH1 [35], NOTCH2 [36], NOTCH3 [37] and NOTCH4). The C-terminal domain is involved in transcriptional activation. The N-terminal domain binds CREBBP/CBP [38] and CDK8 [39]. The gene is reported to be involved in Notch signaling and pre-NOTCH expression and processing pathways. It regulates various genes viz. ID2, NR4A2, FOS, STC1, NR4A1, CREM, ATF3, MAF, CTH, KRT17, PTN, CRYAB, THBS1, CGB3, HAS2 [31], many of them viz. FOS, STC1, NR4A1, MAF, CTH, KRT17, PTN, CRYAB, THBS1, HAS2 [40–49] are known for their association to BRCA. The neurogenic mastermind like N-terminal domain in MAML2 was found to be affected by many deleterious mutations (Fig. 7b). It was interesting to find MAML2 to be downregulated in early stage ER/PR-/HER-2+ samples (log_2__FC=-2.14) (supplementary figure S5b). Being co-activator of NOTCH protein, which is having dual role as oncogene and tumor-suppressor [50], MAML2 might be having tumor-suppressor role in BRCA or might be leading to oncogenesis through some other pathways, but the molecular mechanism is not yet clear. Therefore, it would be interesting to understand role of downregulation of MAML2 in BRCA samples.

#### Early stage TNBC: TTN and interactors

The TTN is most mutated gene followed by SYNE1. The other members of networks were less mutated with somatic mutation frequency (SMF) <= 1%. In TTN, the total mutations were 438, (366-missense, 72-truncating and 4-inframe). In SYNE1, total mutations were 105, (79-missense, 20-truncating, table 6c and supplementary table S6c). The figure 6c shows deleterious mutations in each member. The percent mutation rate for TTN and SYNE1 is significantly high that may make them ideal candidate genes in BRCA.

**Table 6c.** Mutational detail of TTN gene and its interactors in early stage TNBC. The data represents the total unique mutations across the carcinoma samples across four studies (TCGA; Cell 2015, TCGA; Nature 2012, TCGA; PanCancer Atlas and TCGA; Provisional). The cluster harbored several types of mutations as missense, nonsense, fusion, splice, frame-shift, in-frame. The somatic mutation frequency of TTN was found to be 14.4% which is very high as compared to known genes (BRCA1 and BRCA2). Also, SYNE1 showed somatic mutation frequency as 4.4% which is even much greater than BRCA1 (1.7%) and BRCA2 (1.8%).

|  |  |  |  |  |
| --- | --- | --- | --- | --- |
| UNC5B | 09 | Missense, Splice,<br>Frame_shift_deletion | 04 | 0.3 |
| SYNE1 | 105 | Missense, Nonsense,<br>Frame_shift_deletion,<br>Splice, Fusion | 22 | 4.4 |
| NEO1 | 09 | Missense, Splice, Fusion | 05 | 0.6 |
| NTN4 | 03 | Missense, Nonsense | 02 | 0.1 |
| TTC3 | 17 | Missense, Fusion,<br>Nonsense,<br>Frame_shift_deletion | 07 | 0.8 |
| TTN | 438 | Missense, Splice,<br>Nonsense,<br>Frame_shift_deletion,<br>In_frame_deletion,<br>In_frame_insertion,<br>Frame_shift_insertion | 55 | 14.4 |
| MYOM1 | 12 | Missense, Nonsense,<br>Frame_shift_deletion | 07 | 0.7 |
| MYOM2 | 19 | Missense, Nonsense,<br>Frame_shift_deletion | 05 | 1.0 |
| BRCA1 | 42 | Missense, Splice,<br>Nonsense,<br>Frame_shift_deletion,<br>Frame_shift_insertion, | 10 | 1.7 |
|  |  | In_frame_deletion |  |  |
| BRCA2 | 50 | Missense, Splice, Non-sense, Fusion, Frame_shift_deletion, Frame_shift_insertion, In_frame_deletion | 11 | 1.9 |

#### TTN as candidate gene in early stage TNBC

TTN is protein-coding gene associated with myopathy and muscular dystrophy. The gene contains immunoglobulin, kinase, Mex5, N2A and N2b spring domain. It was found to be significantly downregulated (log_2_ FC=-2.9) (supplementary fig. S5c) in early stage TNBC. The immunoglobulin and kinase domains were found to be majorly affected by missense mutations. A very large number of mutations (55 of 438) in TTN were predicted as deleterious, which makes it noteworthy. The gene is involved in actin cytoskeleton signaling, adrenomedullin signaling, integrin signaling, protein kinase A signaling and RhoA signaling. It regulates function of FHL2, ANKRD2, CRYAB, FHL1, PLN, Calm1, ATP2A2, TRIM63, CABLES1 genes [51–56] which might be involved in progression of BRCA. The TTN play role in cell growth and maintain structural integrity of cell. Its molecular function is associated with ankyrin binding and protein kinase activity [57]. The figure 7c shows immunoglobin-like family and fibronectin domains to be significantly enriched in mutations.

#### SYNE1 as candidate gene in early stage TNBC

It is protein-coding gene which encodes spectrin repeat containing protein. The gene is associated with muscular dystrophy and contains KASH (Nuclear envelop localization), calponin homology (CH) and spectrin repeat domain. It is involved in processes as cell death, migration, differentiation and proliferation [58]. The gene regulates glutamate receptor and binds DLGAP1, IFT57, SUN1, TNIK, NDEL1, EMD, LMNA, SYNE1, DISC1, HEY1, DLG4, SUN3, AGAP2, ATP1B4, TUBB2B [31]. The gene is significantly downregulated [59] (log_2_ FC=-1.7) and highly mutated gene in TNBC (supplementary fig. S5d). It was found to have 22 deleterious mutations out of 79 missense mutations affecting CH and spectrin repeat domain (Fig. 7d). The spectrin based domain protects epithelial cells from mechanical stress and is involved in homeostasis of water and salt. The mutation in spectrin domain may lead to its loss of function which results in defective TGF-B signaling and cell cycle deregulation. The loss of this domain is a marker of metastatic cancer cells [60]. The plot shows spectrin/alpha-actinin and spectrin repeat domain got significantly enriched in mutations.

#### Late stage TNBC: DMD, ANK1, ANK2 and interactors

DMD and ANK2 were found to be highly mutated. The DMD is a well-known gene [22] but there are no reports for role of ANK2 in BRCA. ANK2 (SMF=2.0%) directly interacts with DMD (SMF=3.6%) and most mutations in ANK2 and DMD were found to be missense. ANK1 and AHNAK had SMF as 1.5% and 1.4% respectively and other members of cluster were less mutated (SMF<1%). Among all, six were well-known in BRCA while rest (NBPF3, ANK1, NFASC, CHL1, ANK2, CEP 120, GAK2 and CCDC18) are not reported yet. Among these, we found ANK2 as most mutated (table 6d and supplementary table S6d) and the percent mutation rate is higher than BRCA1 and BRCA2. The figure 6d shows deleterious mutation present in each member.

**Table 6d.** Mutational detail of DMD, ANK1, ANK2 and interactors in late stage TNBC. The data represents the total unique mutations across the carcinoma samples across four studies (TCGA; Cell 2015, TCGA; Nature 2012, TCGA; PanCancer Atlas and TCGA; Provisional). The cluster harbored several types of mutations as missense, nonsense, fusion, splice, frame-shift, in-frame. Among the network, ANK2 harbored greater rate of somatic mutation frequency i.e. 2.0%, even greater than BRCA1 and BRCA2.

| Gene name | Total mutations | Mutation type | Mutation showing deleterious impact | Somatic mutation frequency (%) |
| --- | --- | --- | --- | --- |
| CCDC18 | 12 | Missense, Nonsense, Frame_shift_deletion, Splice, Fusion | 01 | 0.5 |
| CEP120 | 11 | Missense, Nonsense, In_frame_deletion | 05 | 0.7 |
| NBPF3 | 06 | Missense, Splice | 03 | 0.2 |
| AHNAK | 47 | Missense, Fusion, Frame_shift_deletion | 0 | 1.4 |
| NBPF1 | 27 | Missense, Splice,<br>Frame_shift_deletion | 03 | 0.9 |
| EHD2 | 06 | Missense, Nonsense,<br>Frame_shift_deletion | 02 | 0.4 |
| NFASC | 21 | Missense,<br>Frame_shift_insertion, Splice,<br>Frame_shift_deletion, Fusion | 08 | 1.0 |
| CHL1 | 08 | Missense, Non-sense | 06 | 0.4 |
| TNS1 | 13 | Missense,<br>Frame_shift_insertion,<br>Frame_shift_deletion,<br>In_frame_deletion | 03 | 0.7 |
| L1CAM | 12 | Missense,<br>Frame_shift_deletion | 07 | 0.8 |
| GALK2 | 06 | Missense, Fusion | 02 | 0.3 |
| DMD | 74 | Missense, Nonsense,<br>Frame_shift_deletion,<br>Frame_shift_insertion, Splice | 29 | 3.6 |
| ANK2 | 40 | Missense, Splice, Non-sense,<br>Frame_shift_deletion | 15 | 2.0 |
| ANK1 | 37 | Missense, Fusion,<br>Frame_shift_deletion,<br>Frame_shift_insertion,<br>Nonsense, Splice | 18 | 1.5 |
| BRCA1 | 42 | Missense, Splice, Nonsense,<br>Frame_shift_deletion,<br>Frame_shift_insertion,<br>In_frame_deletion | 10 | 1.7 |
| BRCA2 | 50 | Missense, Splice, Non-sense,<br>Fusion, Frame_shift_deletion,<br>Frame_shift_insertion,<br>In_frame_deletion | 11 | 1.9 |

#### ANK2 as candidate gene in late stage TNBC

It is protein-coding gene, encoding ankyrin-family of proteins. The binding site in ankyrins facilitates binding of integral membrane proteins with spectrin-actin based membrane cytoskeletal. This linkage maintains integrity of plasma membrane and anchors specific ion channels. They play role in cellular activities e.g. motility and proliferation [61]. The gene is significantly downregulated (log_2_ FC=-2.2) (supplementary fig. S5e) and harbored 40 mutations, out of which 32 were missense. The ankyrin repeat domain and ZU5 domains were mainly affected by missense mutations (Fig. 7e). Ankyrin repeat domains are present in diverse proteins and act as platform for interactions with other proteins. The ankyrin repeat domain binds to miRNA, causing loss in their function and drives proliferation in renal cancer cells [62]. Additionally, the overexpression of ankyrin repeat domain is related with drug resistance in lung cancer [63]. The ZU5 domain is involved in induction of apoptosis and binds to NRAGE domain of UNC5H. The ANK2 regulates ITPR1, ITPR, RYR2, MAPK1, ERK1/2, SPTBN1, focal adhesion kinase, SCN8A and ANK3. Among these, RYR2, MAPK1 and ERK1/2 [64–66] are known to be associated with BRCA. The ANK2 is involved in molecular functions such as protein kinase binding, ATPase binding and structural constituent of cytoskeletal.

#### Genomic variations among the molecular classes

To compare genomic variations (differential-expression and copy-number variations (CNVs)) among three classes in early & late stage BRCA, a circos plot was created (Fig. 8). The outermost track (track 0) is circular ideogram which represents chromosomes. Extending inside, is the heatmap and histograms representing different molecular class of BRCA (Fig. 8a). Corresponding to each of the heatmap plot, there are histograms following same sequence of tracks. In early stage ER/PR+/HER-2-, chromosome 1,7,11,17,19 harbored most of dysregulated genes while in late stage, they are present in chromosome 1,7,11,12,14,15,17,19. In early stage ER/PR-/HER-2+, chromosomes affected were 1,8,12,16,17,19. In TNBC, affected chromosomes were 1,6,10,11,12,16,17,19 in early stages while in late stages, affected chromosomes were 1,2,3,7,10,12,15,16,17, 19,20,22.

**Figure 8.**
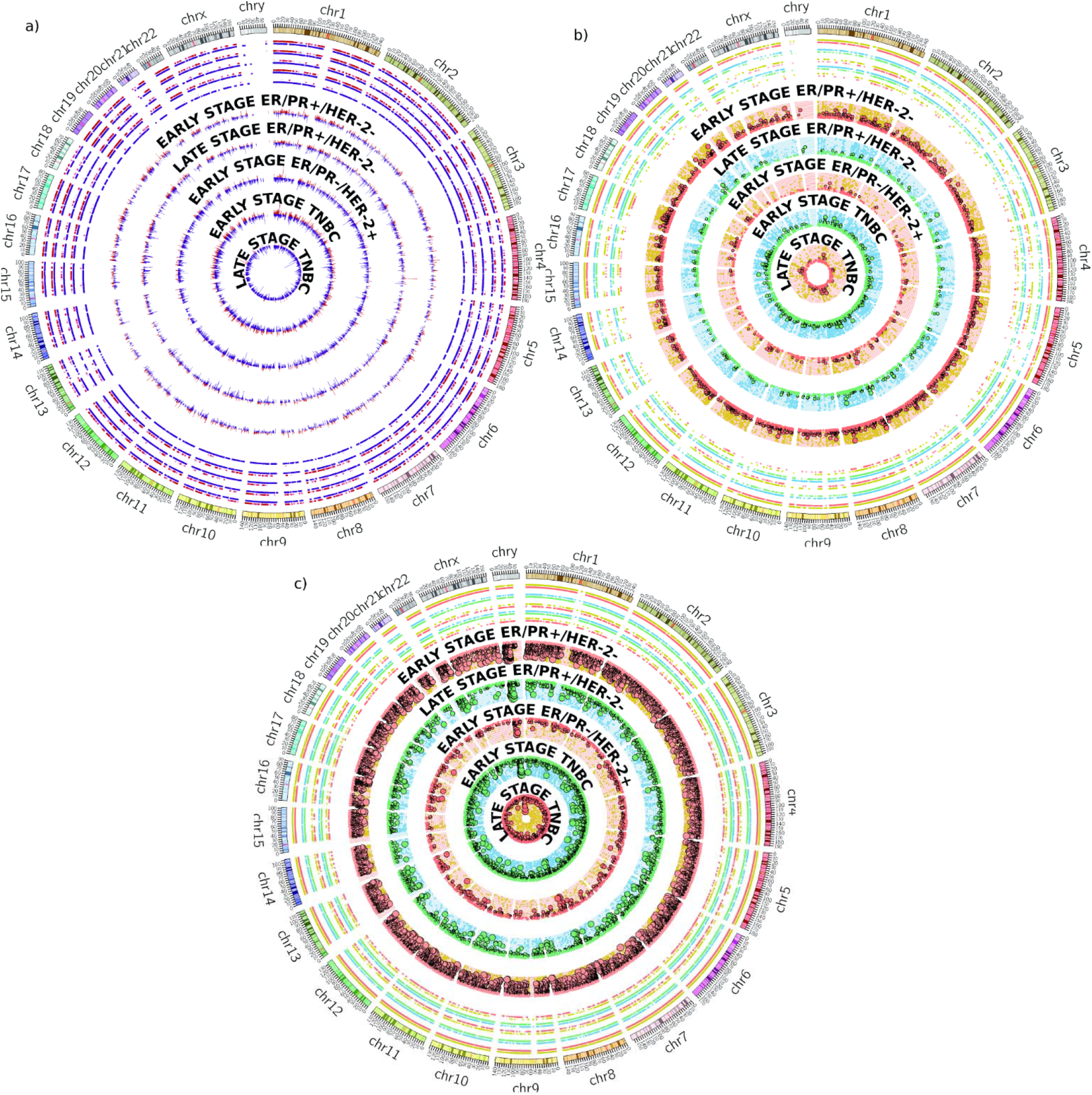
Circos plot representing a) upregulated (red), downregulated (blue) genes, b) gain (pale yellow & blue) and amplification (red & green), c) shallow (pale yellow & blue) and deep deletion (red & green) in three molecular classes of BRCA across whole genome. The outermost track is circular ideogram depicting chromosomes, followed by five heatmap tracks and five histogram tracks. The first track in heatmap represents early stage ER/PR+/HER-2-followed by late stage ER/PR+/HER-2-, early stage ER/PR-/HER-2+, early stage TNBC and late stage TNBC. Same sequence of labelling is followed for histograms tracks as well. The extremely large segment-mean values (>2 or <-2) in b) and c) are outlined in black.

The CNV data was obtained from TCGA includes amplification (Fig. 8b) and deletion (Fig. 8c) events in BRCA. The segment mean value from 0.5-1 represents gain (pale-yellow and blue) while >1 represents amplification (red and green). The CNVs are shown as scatter plots and for extremely large values (>2), black outlines have been put around glyphs. In early stage ER/PR+/HER-2-, amplification enriched chromosomes 7,8,11,12,17 while in late stage, chromosome 1,8,17,20 were enriched. In early stage ER/PR-/HER-2+, chromosome 1,8,11,17 were enriched in amplifications and in TNBC, chromosome 1,5,6,8,16,17 were affected in early stage while chromosome 8,16 were affected in late stage. The segment mean value from −0.5 to −1 represents shallow deletion (pale yellow and blue) while <-1 deep deletion (red and green). Extreme values (<-2) are outlined in black. The plot depicts that most of the chromosomes were enriched in deep deletions events in early and late stages.

#### Survival analysis

The survival analysis aimed to evaluate overall survival of BRCA patients having mutations in NLGN3, MAML2, SYNE1, TTN and ANK2 genes. It can reveal the prognostic value of these genes. The Kalpan-Meier curves for NLGN3, MAML2, SYNE1, TTN, ANK2, BRCA1 and BRCA2 are shown in supplementary figures S6a, S6b, S6c, S6d, S6e, S6f and S6g respectively. The X-axis represents overall survival of cases with and without alteration in gene and Y-axis represents survival in months. The curve represents that overall survival of cases with alteration in NLGN3, MAML2, SYNE1 and TTN gene is less than the cases without alteration in these genes [27]. The log-rank test p-value for NLGN3, MAML2, TTN, SYNE1 and ANK2 was estimated as 0.951, 0.071, 0.7, 0.712 and 0.298 respectively. In case of BRCA1 and BRCA2, p-value was estimated to be 0.818 and 0.988. The log-rank test p-value indicates that there is no significant difference between the overall survival in both groups. This indicates that the mutations in these genes has no significant impact on survival.

#### Experimental validation

Validation of predictions is quintessential part of scientific research. In the current studies to validate the predictions, one gene from each of the subnetwork was picked, thus four genes viz. NLGN3, ANK2, MAML2 and SYNE1 were selected. The transcript level of selected genes in different breast cancer cell lines MCF7, T47D, MDA-MB-231 was determined. The transcript levels were compared with normal breast epithelial cell line MCF10A using qRT-PCR. To determine the difference in expression of these genes in cancer versus normal cell-line, comparative CT method was applied taking GAPDH as endogenous reference gene. The amplification of GAPDH was done earlier than other candidate genes as revealed by CT value, indicating that the genes are down-regulated in cancer cell lines. The expression of NLGN3 was 0.12, 0.15, 0.30 fold downregulated in MCF7, T47D and MDA-MB-231 respectively. The expression of MAML2 was 0.16, 0.33 and 0.29-fold (downregulated) in MCF7, T47D and MDA-MB-231 respectively. The expression of SYNE1 was 0.08 and 0.17 fold (downregulated) in MCF7 and MDA-MB-231 respectively and 1.44-fold upregulated in T47D. The expression of ANK2 was 0.06, 0.27and 0.24-fold (downregulated) in MCF7 and MDA-MB-231 respectively (Fig. 9a,b,c and d).

**Figure 9.**
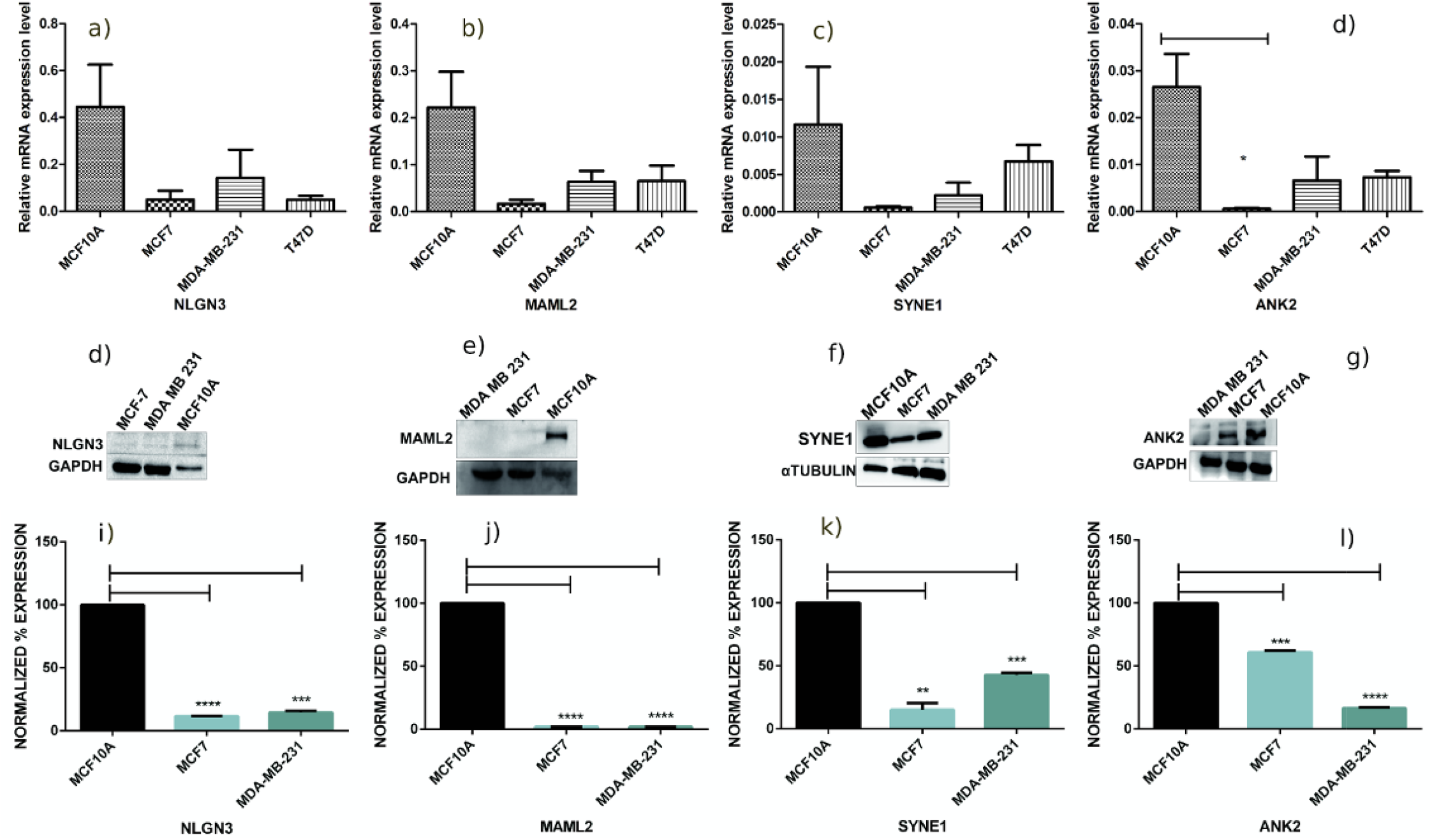
Real-Time PCR and western blot analysis of selected genes viz. NLGN3, MAML2, SYNE1 and ANK2. Figure shows relative mRNA expression level of a) NLGN3 b) MAML2 c) SYNE1 d) ANK2 in MCF10A, MCF7, MDA-MBA-231 and T47D cell lines. The GAPDH was used as control. The statistical analysis was done by unpaired student’s t-test. Western blotting data of e) NLGN3, f) MAML2, g) SYNE1 and h) ANK2 is depicted with GAPDH and alpha-tubulin expression level. The blots were quantified by densitometry, normalized and subjected to statistical analysis and figures are shown in i), j), k) and l) respectively for NLGN3, MAML2, SYNE1 and ANK2.

#### Western blot analysis

Western blotting exhibited the protein expression of NLGN3, MAML2, SYNE1 and ANK2 is decreased in cancer cell lines (MCF7 and MDA-MB-231) as compared to control cell line (MCF10A) confirming our earlier observations. The blots of NLGN3, MAML2, SYNE1 and ANK2 are represented in figure 8e, f, g, and h respectively. The unpaired student’s t-test showed the decrease in protein expression of NLGN3, MAML2, SYNE1 and ANK2 was significant in both the cancer cell lines e.g. MCF7 and MDA-MB-231 (p-value<0.01) when compared with control MCF10A (Fig. 9i-l).

#### Immunohistochemistry

##### NLGN3

The mean expression of neurolgin3 was found to be 7.91, 7.86, 8.5, 6.0 and 13.2 in early stage ER/PR+/HER-2-, late stage ER/PR+/HER-2-, early stage TNBC, late stage TNBC and NATs samples respectively, which reveals that the neurolgin3 is downregulated in each of the four categories. The Mann-Whitney test revealed that the significant downregulation of NLGN3 in early stage ER/PR+/HER-2- (p-value=0.0162) and early stage TNBC (p-value=0.0328) (fig.10a, 10d). This corroborates with our predictions in early stage ER/PR+/HER-2-class. The grade-wise comparison showed that the NLGN3 is significantly downregulated in grade 2 (p-value=0.0137) and grade 3 (p-value=0.0123) (fig.10g, 10j).

**Figure 10.**
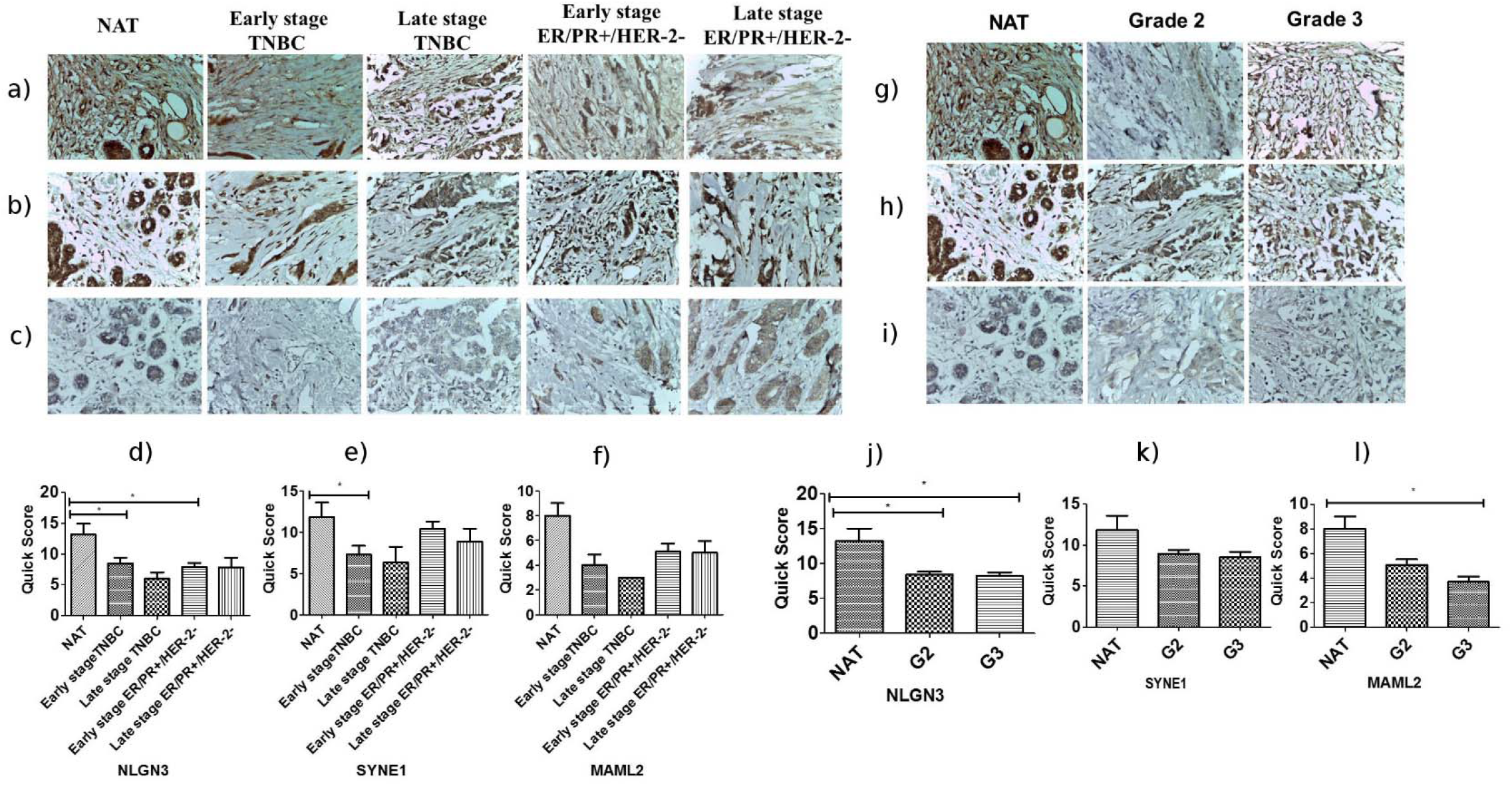
Figure showing the immunohistochemically stained TMA sections for a) NLGN3 b) SYNE1 and c) MAML2 in NAT, early stage TNBC, late stage TNBC, early stage ER/PR+/HER-2- and late stage ER/PR+/HER-2-. d) NLGN3 expression is significantly decreased in early stage ER/PR+/HER-2-(p-value=0.0162) and early stage TNBC (p-value=0.0328). e) SYNE1 expression is significantly downregulated in early stage TNBC (p-value=0.0410). f) Downregulated expression of MAML2 was found in all 4 categories but was not statistically significant. Right panel shows the immunohistochemically stained TMA sections for g) NLGN3 h) SYNE1 and i) MAML2 in NAT, grade 2 and grade 3 tumor. j) The expression of NLGN3 is significantly decreased in grade 2 (p-value=0.0137) and grade 3 (p-value=0.0123) compared to NAT. k) There was a decrease is SYNE1 expression in grade 2 and 3 tumor samples but it was statistically insignificant. l) MAML2 expression is significantly decreased in grade 3 (p-value=0.017).

##### SYNE1

The tissue expression of SYNE1 is decreased in each of the four categories. The mean expression was found to be 10.41, 8.86, 7.25, 6.33 and 11.83 in early stage ER/PR+/HER-2-, late stage ER/PR+/HER-2-, early stage TNBC, late stage TNBC and NATs respectively. The Mann-Whitney test revealed that the significant decrease in tissue expression of SYNE1 in early stage TNBC class (p-value=0.0410) whereas in other classes, the decreased expression was not statistically significant (fig.10b, 10e). The decrease in tissue expression from early stage TNBC to early stage ER/PR+/HER-2-was statistically significant (p-value=0.0306). The tissue expression of SYNE1 is also decreased in grade 2 and 3 tumors but was not statistically significant (fig. 10h, 10k).

##### MAML2

The mean expression of MAML2 was found to be 5.11, 5, 4, 3, 8 in early stage ER/PR+/HER-2-, late stage ER/PR+/HER-2-, early stage TNBC, late stage TNBC and NATs respectively. The tissue expression of MAML2 is downregulated in early stage ER/PR+/HER-2- and early stage TNBC but was not statistically significant (fig.10c, 10f). Due to non-availability of HER-2+ samples in TMA slides, tissue expression could not be checked in it. The grade-wise analysis showed that the MAML2 is significantly downregulated in grade 3 (p-value=0.017). The tissue expression was to decrease progressively from grade 2 to grade 3 (p-value=0.047) (fig. 10i, 10l).

##### ANK2

The immunohistochemistry was performed but no significant staining was found in NAT and breast carcinoma samples.

## Discussion

BRCA is the biggest health problem in females. It remains undetected until the advanced stages which then becomes the primary reason for high mortality rates. The development of carcinoma can be influenced by the genetic as well as environmental variations which affects the cellular and functional changes. The major research problems associated with BRCA include recognition at early stage, anticipation of re-occurrence and tackling the emergence of drug resistance. Many efforts are going on worldwide to tackle the menace of invasive breast carcinoma. The AACR report-2018 summarizes the worldwide statistics, diagnosis options and treatments available to the breast cancer patients.

Among the three molecular classes, TNBC is considered the most aggressive form. The five-year survival rate is lower with TNBC than other breast cancer types perhaps due to lack of molecularly targeted therapies. Therefore, efforts are being done in this direction e.g. a ten gene expression signature has been reported to be associated with TNBC, in particular for Mexican patients [67]. Also, in TNBC different dysregulated non-coding RNAs were analysed [68] and molecular pathways were identified [69]. The overall and disease-free survival was related to certain gene expression in TNBC [70, 71].

The genomics differences between African and Caucasian American women were studied and 20 genes were identified that segregated two classes [72]. ITGA11 and Jab1 were identified as biomarker for breast cancer [73]. The mutations in important loci as BRCA1, BRCA2, PTEN, ATM, TP53, CHEK2, PPM1D, CDH1, MLH1, MRE11, MSH2, MSH6, MUTYH, NBN, PMS1, PMS2, BRIP1, RAD50, RAD51C, STK11 and BARD1 were related with risk of breast cancer [74]. The relation between somatic mutations and prognosis was determined and mutations in PIK3R1 and DDR1 were found to be associated with poor outcomes in hormone receptor positive breast cancer [75]. The cancer genes modules containing low-frequency genes were identified in breast cancer [76].

This is the first study to identify subnetworks of genes in three molecular classes (ER/PR+/HER- 2-, ER/PR-/HER-2+ and TNBC) in different clinical stages based on gene dysregulation and mutations.

This study reports five genes viz. neuroligin3 (NLGN3), mastermind like transcriptional coactivator 2 (MAML2), titin (TTN), ankyrin2 (ANK2) and spectrin repeat containing nuclear envelop protein 1 (SYNE1) to be significantly associated with distinct molecular classes of BRCA (ER/PR+/HER-2-, ER/PR-/HER-2+ and TNBC). The genes are involved in important processes such as chemotaxis and axon guidance, notch binding, cell adhesion molecule binding etc. They are central genes in the protein-protein-interaction (PPI) network indicating they can have important regulatory roles. The alteration in their expression levels and protein function may greatly impact the expression and functions of downstream genes, which initiate and accelerate the process of oncogenesis. The identified genes were found to be involved in different biological pathways as TR/RXR activation pathway, Notch signalling pathway, regulation of EMT, Netrin signalling, RhoA signalling, axonal guidance signaling pathways and galactose degradation I pathway. These pathways have important roles in cellular processes such as proliferation, differentiation, migration, survival and predispose oncogenesis.

In early stage, pathways as interferon signaling, tryptophan degradation III, granulocyte adhesion & diapedesis and catecholamine biosynthesis were found in ER/PR+/HER-2- (p-value< 0.010), pathways as RAR activation, adipogenesis, role of JAK1, JAK2 and TYK2 in interferon signaling, TGF-β signaling and STAT3 signaling (p-value < 0.014) intricated in ER/PR-/HER-2+ and pathways as IL-1/IL-8 signaling, TNFR1/TNFR2 signaling, NER, TWEAK signaling and relaxin signaling (p-value < 0.005) were found to be associated with TNBC. In late stage, calcium signaling, cholesterol biosynthesis, FXR/RXR activation and AMPK signaling pathways were (p-value < 0.010) found to be associated with ER/PR/+/HER-2- and E1F2 signaling, mTOR signaling, renin-angiotensin, SAPK/JNK signaling and IL-10 signaling pathways (p-value < 0.014) were found in TNBC.

The findings were further validated using qRT-PCR, western blotting and immunohistochemistry. The genes as NLGN3, MAML2 and ANK2 were found to be downregulated in MCF-7, T47D and MDA-MB-231 cell-lines. SYNE1 was found to be downregulated in MCF-7 and MDA-MB-231. Out of four genes, ANK2 was found to be significantly downregulated at mRNA level in MCF7 cell-line (p-value 0.021). The western blotting experiment also showed the protein expression of NLGN3, MAML2, SYNE1 and ANK2 was significantly downregulated in MCF7 and MDA-MB-231 as compared to MCF10A cell lines which is strongly in-line with our findings. The immunohistochemistry also showed the decreased in expression level of NLGN3, MAML2 and SYNE1 in tissue samples.

The present study revealed that the genes viz. NLGN3, MAML2, TTN, SYNE1 and ANK2 are significantly correlated with BRCA. They interact with numerous other genes associated with cell proliferation, survival, migration, metastasis. The genes initiate and stimulates proliferation of cancer cells by impeding the functions of downstream genes and disrupting normal cellular processes.

The results suggest that candidate genes may serve as useful biomarker for different molecular classes of BRCA. The subnetworks containing gene components can be targeted for developing novel therapeutic regimen. We believe that our comprehensive study can contribute in diagnosis and therapeutics.

## Material and methods

The different molecular classes of BRCA have been analyzed for genes that are differentially expressed in early and late stages, dysregulated biological pathways, and cancer progression using integrated computational analysis. The flowchart of the methodology is shown in Fig. 1

### Data Collection

The transcript expression data for BRCA tissue and normal adjacent tissue (NAT) was obtained from cancer RNA-seq nexus (CRN) **(JIAN *et al*.**,) [77]. The data contained transcripts expression values (total 71,361 (coding and non-coding)) in TPM (transcript-per-million basepairs) of 1097 tumour & 114 NAT samples. The non-coding (6,287) transcripts were removed and the data was divided into three molecular classes a) ER/PR+/HER-2- [tumour samples (362) vs NAT (37)], b) ER/PR-/HER-2+ [tumour (36) vs NAT (04)] and c) Triple negative breast cancer (TNBC) [tumour (113) vs NAT(10)]. Under each molecular class, data was further divided into clinical stages (stage I-IV as per standard TCGA classification). The number of tumour and NAT samples in each class is shown in table 1. However, due to small sample size in each stage, we classified data into two groups i.e. early (I, II) and late (III, IV) stage for analysis purpose. The transcripts with significant p-value (<0.05) were kept for further analysis. The mean expression of transcripts that belong to a gene was considered as expression for that gene.

### Differential expression study

The fold change was calculated using TPM values in tumour and NAT. The genes with a log_2_ FC greater than or less than 1 and p-value<0.05 were considered up or downregulated respectively [78]. The differential expression pattern was studied in the early and late stages in each of the three class. Further, due to non-availability of NAT sample in late ER/PR-/HER-2+ class, it was removed from the study. Finally, five classes (i) Early stage ER/PR+/HER-2- (ii) Late stage ER/PR+/HER-2- (iii) Early stage ER/PR-/HER-2+ (iv) Early stage TNBC and (v) Late stage TNBC were analyzed in detail. The biological pathways in each class were also analyzed to check similarity and differences in terms of affected pathways in different classes.

### Clustering of genes into subnetworks

The genes were clustered based on differential expression, protein-protein interaction (PPI) and mutational frequency using Hotnet2 algorithm. The hotnet2 is an algorithm to find out mutated clusters and overcomes limitations with existing algorithms as MEMo, HotNet etc. These clusters can have co-occurring mutations across cancer samples and their component genes may work together with upstream/downstream genes to drive biological pathway(s). The purpose was to uncover modules, pathways and processes which are getting affected in different classes of BRCA. The mutation data for BRCA was collected from ICGC [79] and subnetworks were created using Hotnet2 algorithm (supplementary data 1).

### Survival analysis

Survival analysis is a statistical method for data analysis in which outcome is any event such as recurrence of disease, disease-free period, death etc. The Kalpan-Meier survival curves estimated the overall survival of BRCA patients with or without alteration in selected genes and have been taken from cBioPortal.

### Pathway analysis

Pathway analysis was performed using IPA (Ingenuity-Pathway-Analysis) using p-value cutoff of 0.05 to uncover functional role of various modules (subnetworks).

### Quantitative-real-time polymerase-chain-reaction (qRT-PCR)

Breast cancer cell lines MCF-7, T47D and MDA-MB-231 were purchased from National Centre for Cell Sciences (NCCS), Pune, India and total RNA was isolated from MCF7, T47D, MDA MB-231 and MCF10A cells using Tri-reagent (Sigma-Aldrich). Reverse transcription PCR and qRT-PCR was performed using primers of NLGN3, MAML2, SYNE1 and ANK2. The detailed methodology and primer sequences are given in supplementary data 2. GAPDH was taken as an internal control and ΔΔCT values were calculated. The statistical analysis was performed using GraphPad Prism, which includes unpaired student’s t-test for estimation of statistical significance.

### Western blot analysis

The whole cell lysate of MCF7, MDA-MB-231 and MCF10A were prepared using RIPA buffer. The samples were run in 10% SDS-PAGE gel, transferred on PVDF membrane (Millipore) and blocked with 5% (w/v) non-fat milk (Sigma). Blots were then incubated with primary antibody (MAML2, ANK2, SYNE1, NLGN3) overnight. The blots were then subjected to chemi- lumenescent detection reagent for visualization and the bands were detected by using Gel Doc™ XR _+_ Imager. Densitometry analyses of protein bands were done by using ImageJ. The detailed methodology is given in supplementary data 3.

### Immunohistochemistry

The tissue microarray slides (TMA) were purchased from US Biomax for each of the genes (NLGN3, MAML2, SYNE1 and ANK2). Each slide contained hundred samples of invasive ductal carcinoma and nine samples of normal adjacent tissue. The samples belonged to different molecular classes, clinical stages, age and grade. Corresponding to our analysis, we have divided the samples into four categories i.e. Early stage ER/PR+/HER-2-, Late stage ER/PR+/HER-2-, Early stage TNBC and late stage TNBC. The number of samples in early stage ER/PR+/HER-2-, late stage ER/PR+/HER-2-, early stage TNBC and late stage TNBC were 23, 7, 13 and 3 respectively. There are 57 samples of grade 2 and 42 samples are of grade 3. The age-wise distribution was as follows: 23 (<= 40 years), 44 (41-50 years), 22 (51-60 years), 11 (>60 years).

Four slides were stained by anti-MAML2, anti-Ank2, anti Syne-1 and anti-NLGN3 antibodies separately at 1:50 dilution and were further processed using ABC system (Vector Laboratories, Bulingame, CA, USA) as described previously [80]. The images were captured under Leica microscope using LAS EZ software version 2.1.0 at 40x magnification. The tissue spots were scored by pathologist in a blinded experiment. The score for intensity of staining and percentage positive cells were given as per Allred scoring system. The staining intensity was scored as: 0 for negative, 1 for weak, 2 for intermediate and 3 for strong. The percentage positive cells were scored between 0%-100%.: 0 for no cells, 1 for <1% cells, 2 for 1-10%, 3 for 11-33%, 4 for 34-66% and 5 for 67-100%. Quick score for each sample was calculated by multiplying intensity score by score for percentage positive cells. The statistical analysis was done with Quick score and graph was plotted.

## Conclusion

Our integrated bioinformatics approach has enabled the identification of dysregulated genes in ER/PR+/HER-2-, ER/PR-/HER-2+ and TNBC in early and late stage of BRCA. During analysis, we found some interesting pattern of dysregulation i.e. the genes which were upregulated in one molecular class, were downregulated in other class and vice-versa. Additionally, some genes were found to be class specific. This difference might be due to difference in expression patterns of hormone receptors among the molecular classes. Later, the significantly mutated subnetworks containing dysregulated, mutated and driver genes were identified in different molecular classes of BRCA which gave insight about their biological functions. We believe that the dysregulated genes can serve as biomarkers for early detection of the class and stage of BRCA. Our approach seeks to maximize the use of datasets and techniques to understand the role of coding genes in disease pathogenesis, prognosis, development of diagnostic tools and therapeutics. We hope that identifying genes and their subnetworks will contribute to drug design and discovery.

## Declarations

### Authors’ contributions

The study was devised by SA, AD and SKK. The experimental validation work was done by SN under the supervision of SKM. The IHC was done by AKA. The useful discussions were done with PK. The manuscript was written by SA, AD & SKK and reviewed by AD, SKM and SKK.

## Acknowledgements

SA wants to acknowledge Department of Science and Technology (DST), Govt. of India, for providing research fellowship under DST-INSPIRE fellowship program.

## Funding

The funding is given by DST-INSPIRE, New Delhi.

## Notes

### Competing Interest Statement

The authors have declared no competing interest.

