## Supplementary tables and figures for "Stagewise identification of significantly mutated and dysregulated gene clusters in major molecular classes of breast invasive carcinoma": Suplementary_figures.pdf

#### Supplementary fig. S1

### TR/RXR Activation pathway

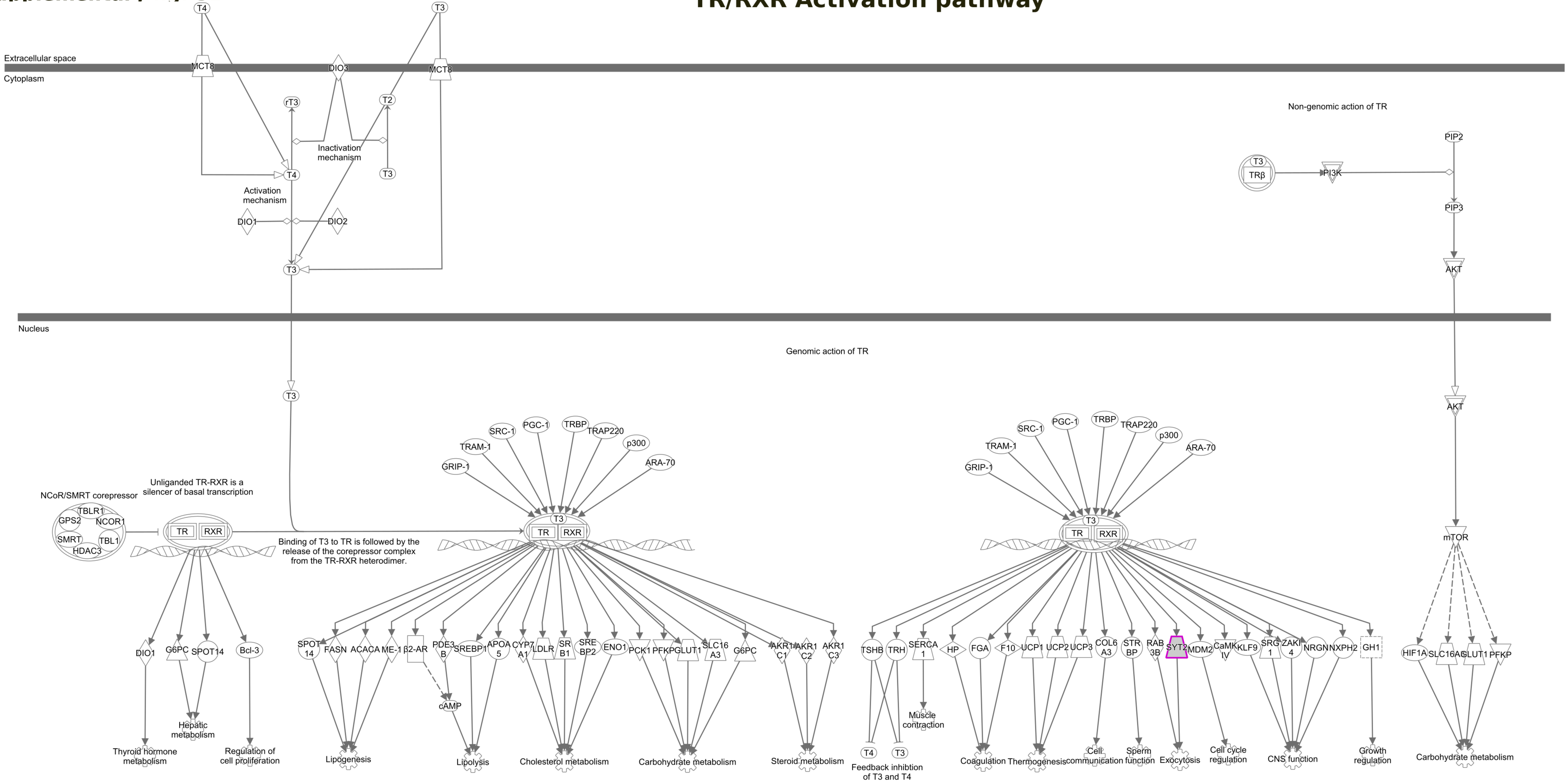


© 2000-2019 QIAGEN. All rights reserved.

Supplementary fig. 2a

### NOTCH Signaling

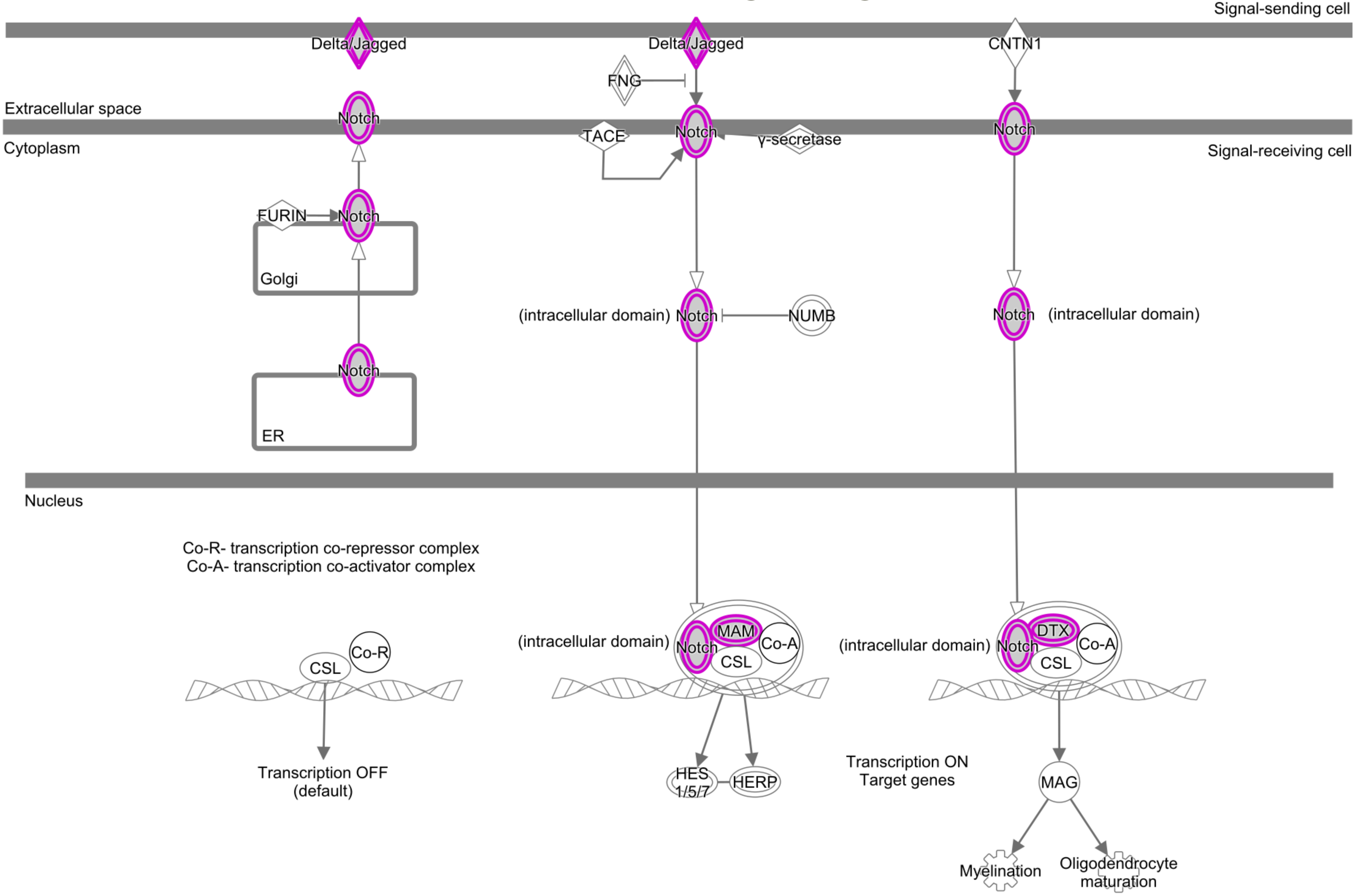


© 2000-2019 QIAGEN. All rights reserved.

#### Supplementary fig. S2b

### TH1 and TH2 Activation pathway

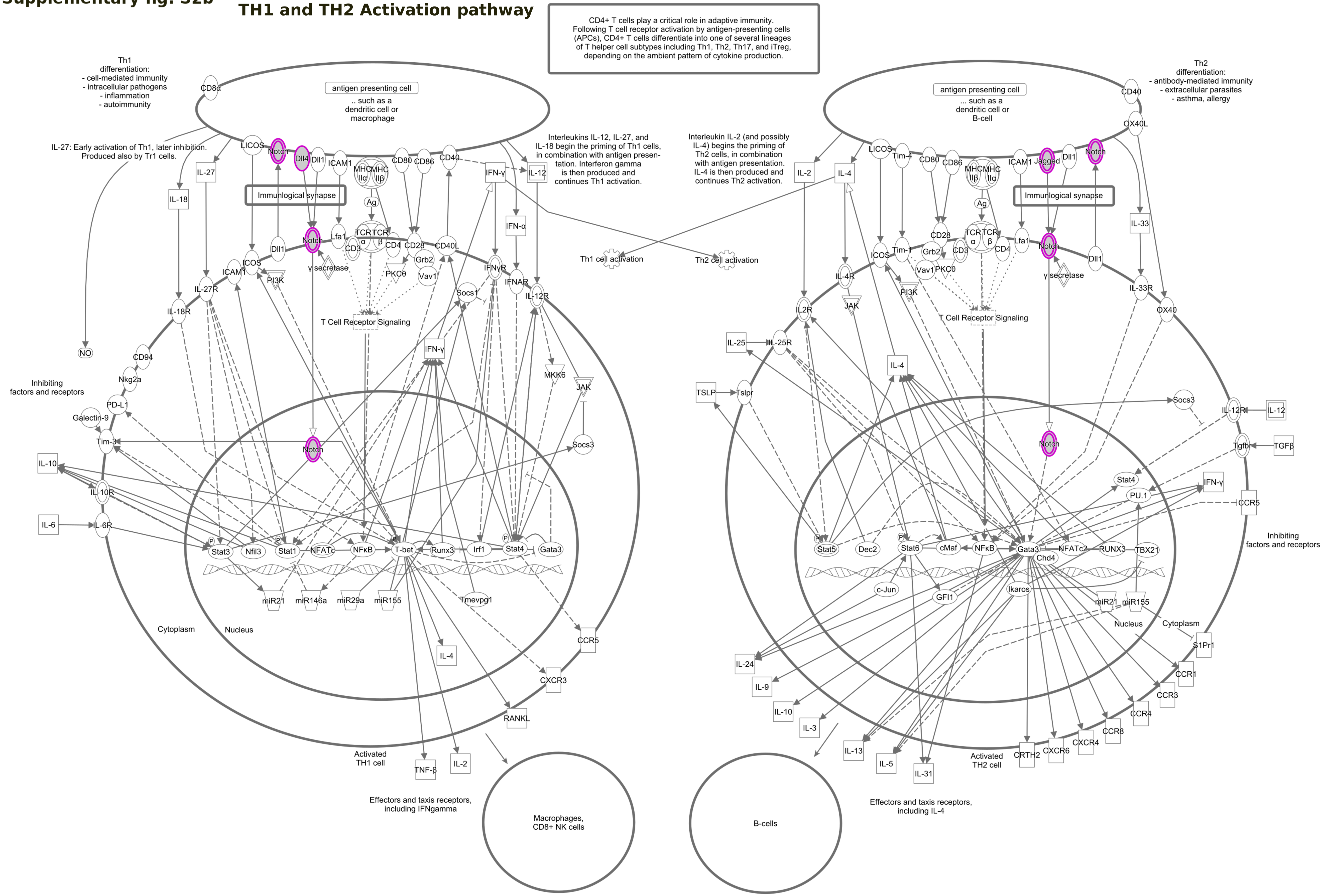


© 2000-2019 QIAGEN. All rights reserved.

#### Supplementary fig. S2c

### Regulation of the Epithelial Mesenchymal Transition pathway

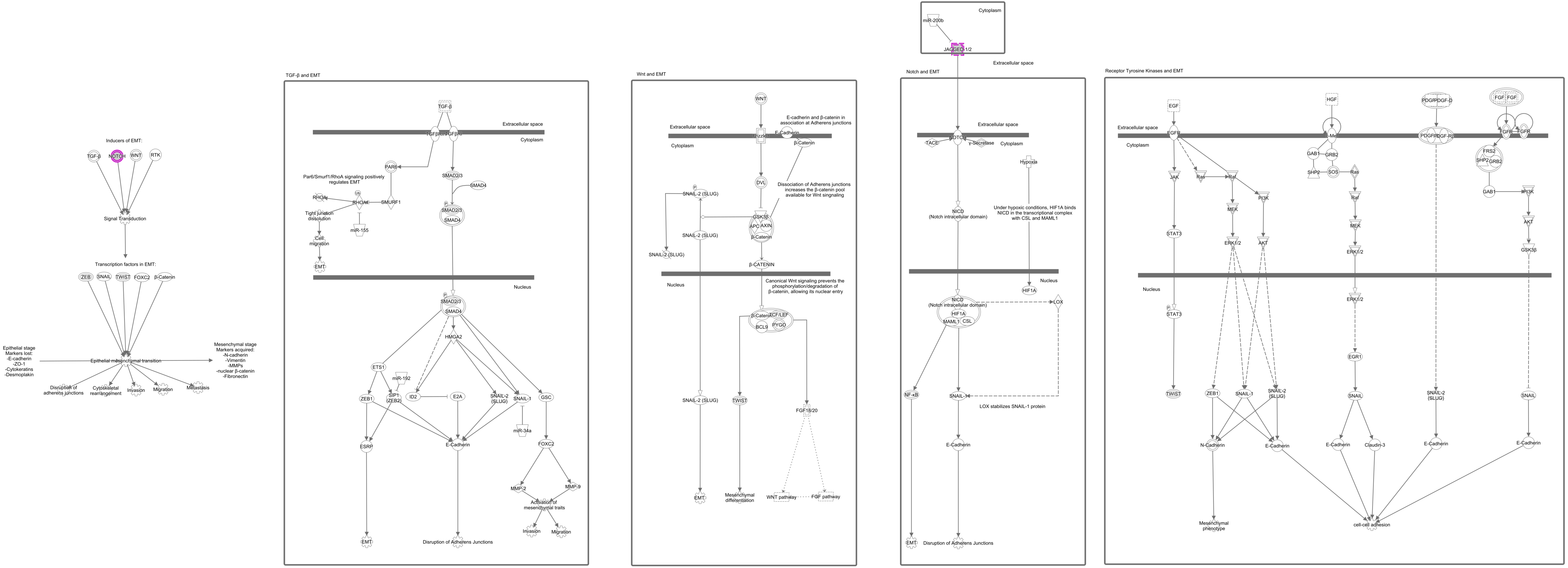


© 2000-2019 QIAGEN. All rights reserved.

**Supplementary fig. 3a**

### Netrin Signaling

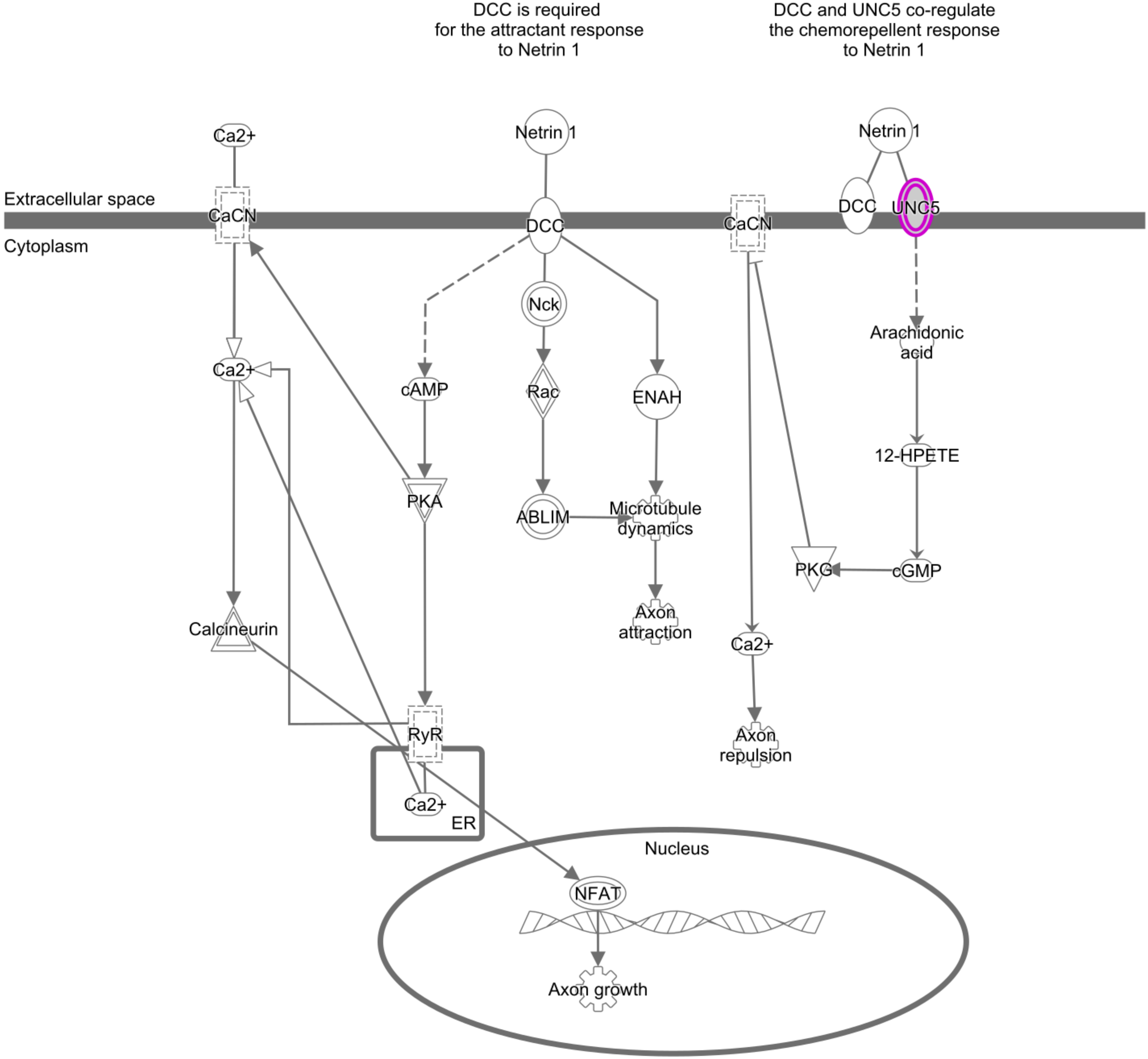


© 2000-2019 QIAGEN. All rights reserved.

### Supplementary fig. S3b

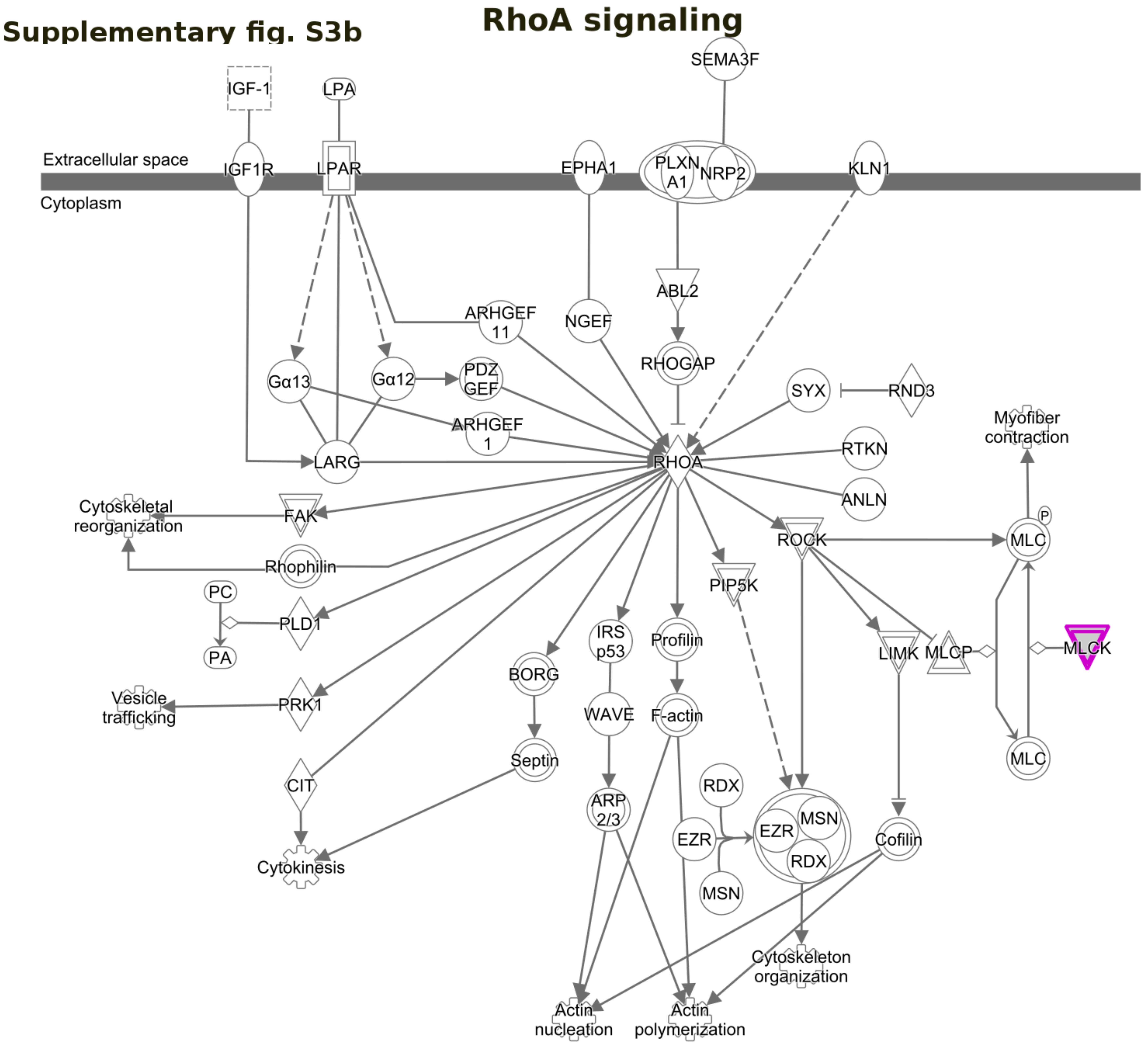


© 2000-2019 QIAGEN. All rights reserved.

**Supplementary fig. S3c**

### Axonal guidance signaling

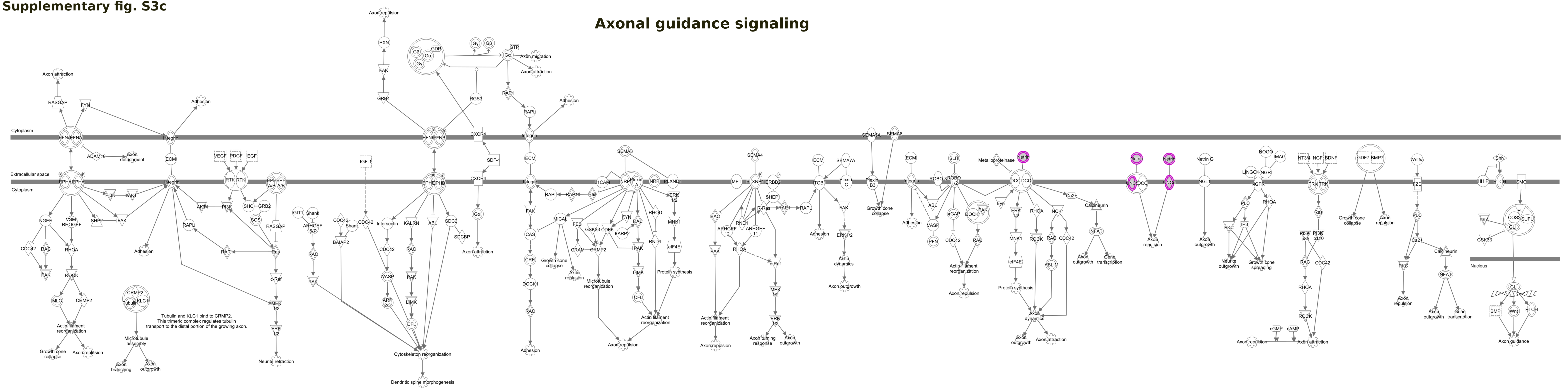


© 2000-2019 QIAGEN. All rights reserved.

### Supplementary fig. S4a

#### Galactose degradation I(Leloir pathway)

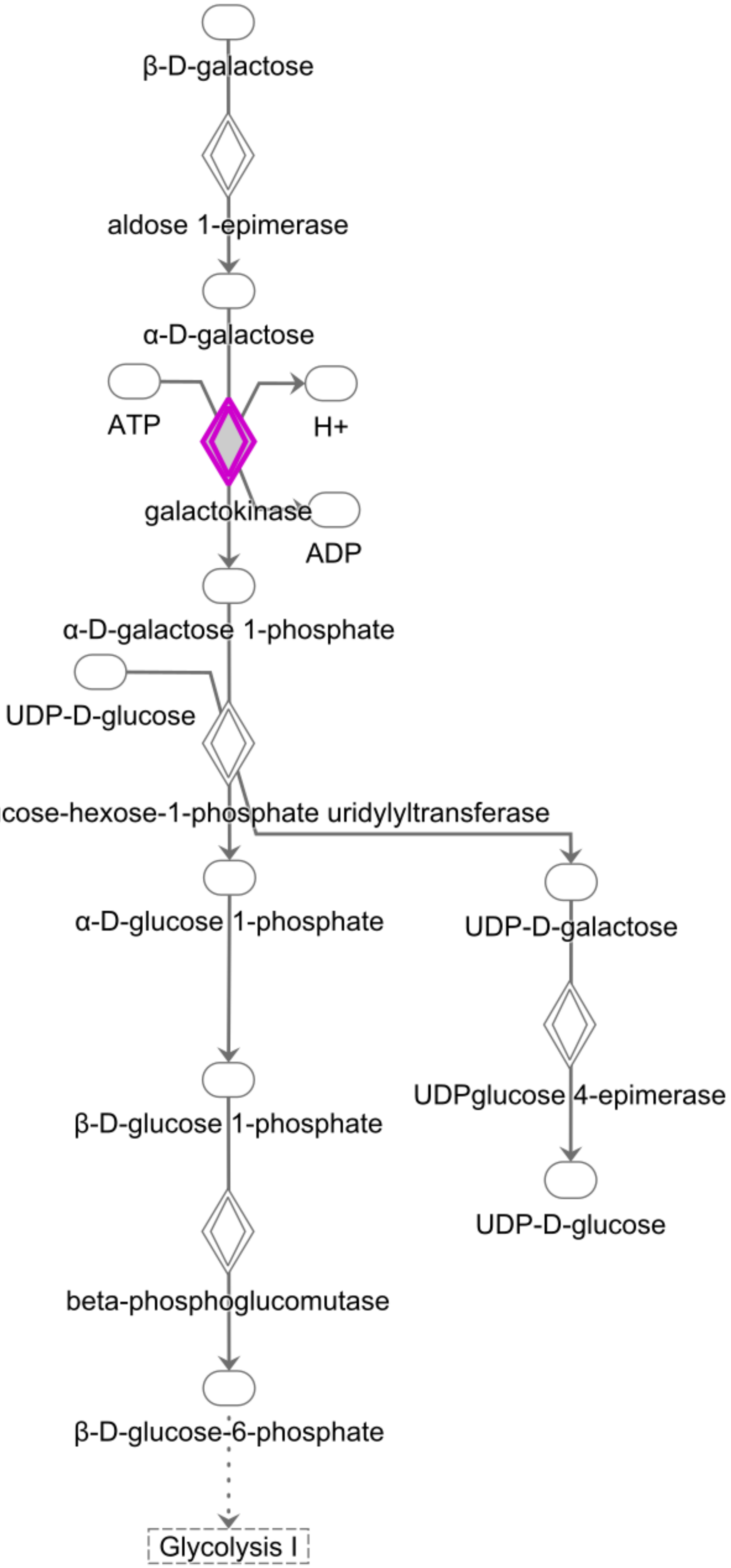


Adapted from HumanCyc.© 2000-2019 QIAGEN. All rights reserved.

Supplementary fig. S4b

### Colanic Acid Building Blocks Biosynthesis

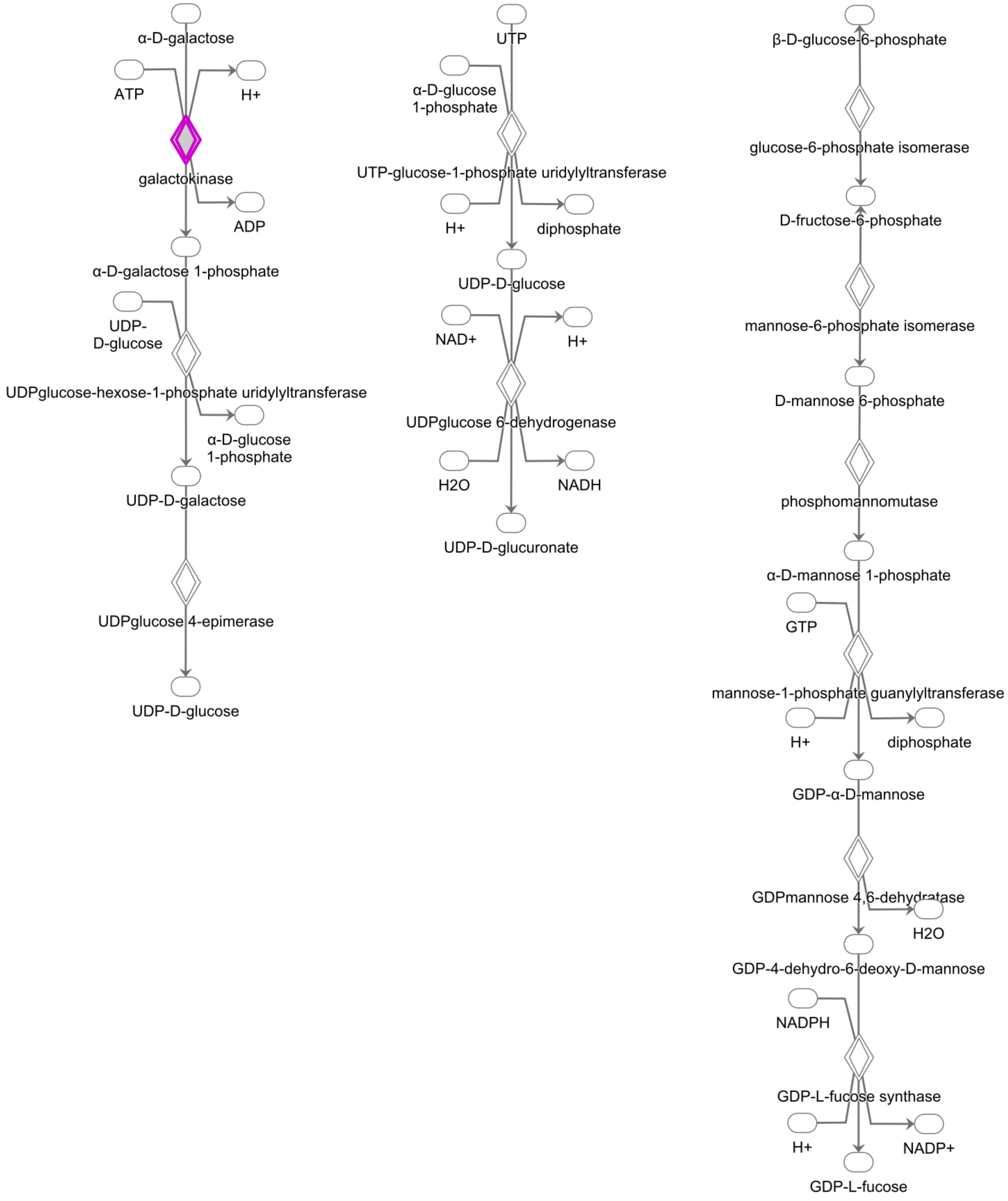


Adapted from HumanCyc.© 2000-2019 QIAGEN. All rights reserved.

Supplementary fig. S4c

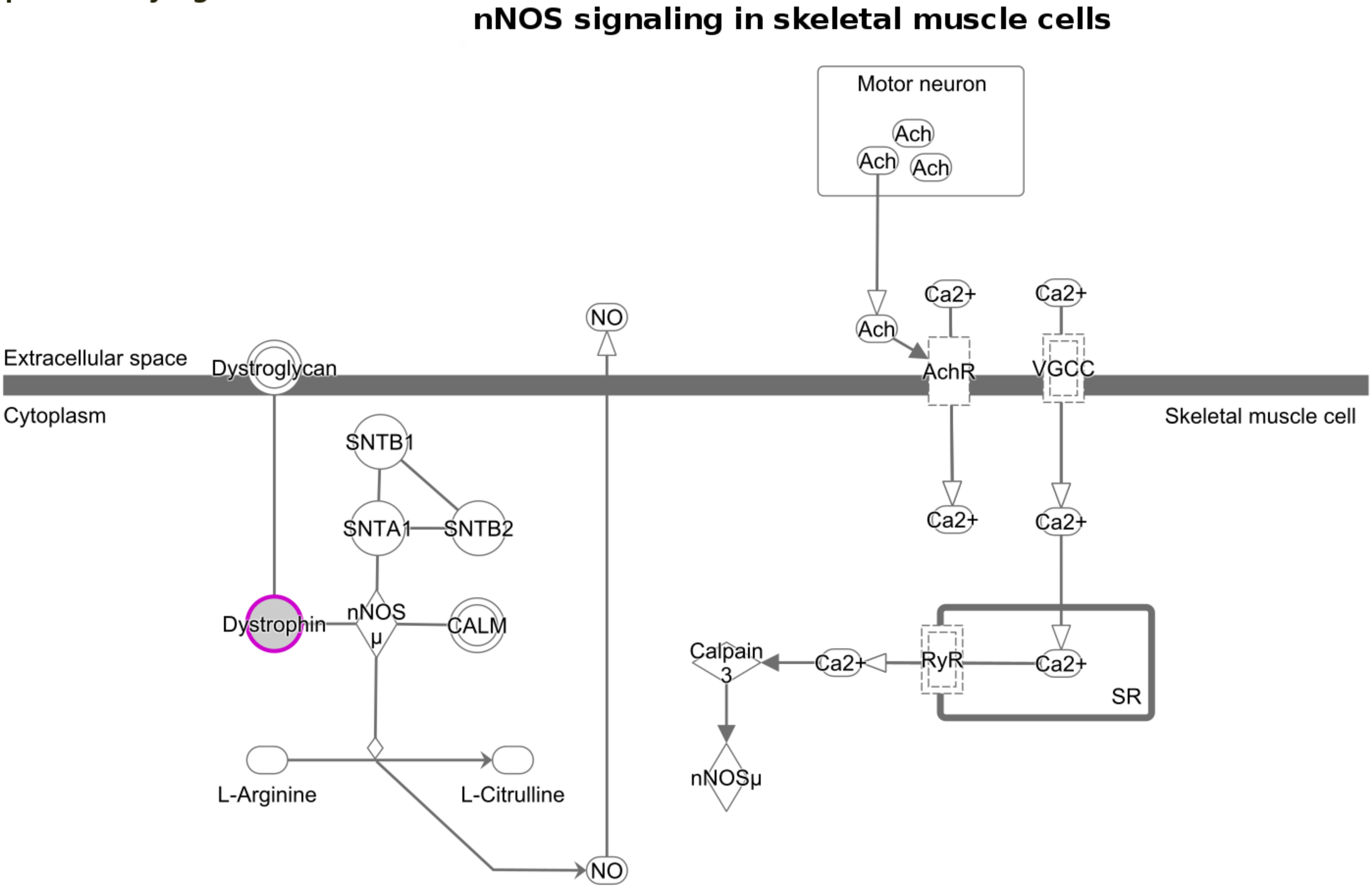


© 2000-2019 QIAGEN. All rights reserved.

#### Supplementary fig. S5

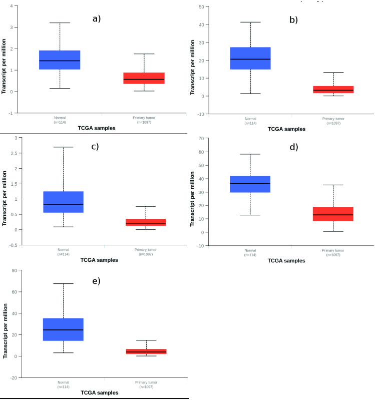

#### Supplementary fig. S6

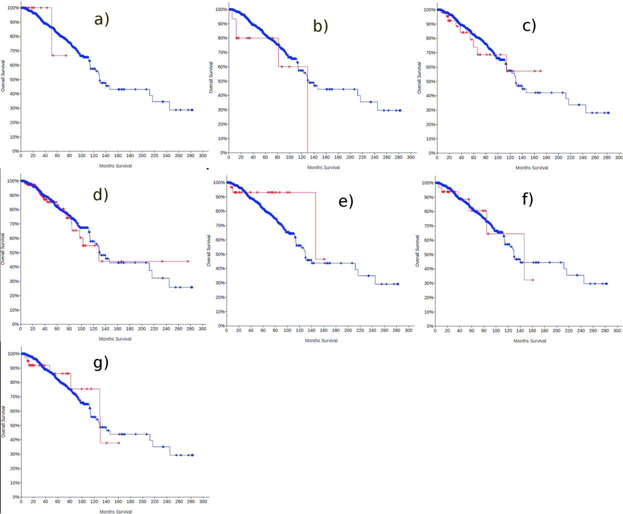
