## Supplementary tables and figures for "Stagewise identification of significantly mutated and dysregulated gene clusters in major molecular classes of breast invasive carcinoma": supplementary_data.docx

1. **Creation of gene clusters using hotnet2**

The Hotnet2 algorithm assigns a heat score to each node (gene) in PPI network based on its frequency of mutation. Hotnet2 finds significantly mutated subnetworks that send and receives a significant amount of heat. It allows nodes to retain some amount of their own heat in the process of heat transfer to neighboring nodes. The amount of heat retained by each node is determined by an insulating parameter β. The β is a point in inflection point diagram before which heat from a protein starts diffusing to neighbour of neighbour instead of direct neighbour. For selection of β, 20 different matrices were created with beta ranging from 0.05 to 1.0. The edge list and index gene files were created using PPI data. The influence matrix file was generated for real (HIPPIE) network. Then, 100 permuted networks were generated for real network. For each of permuted network, influence matrix was created using calculated beta value (0.47). Finally, a heat score is assigned to each of the gene. The delta (edge weight parameter) was used to generate hot clusters of significantly mutated genes. The delta parameters ensure that the hot clusters are found in the real network and not in the permuted ones. The clusters were created for each class in early and late stage. The top significant clusters for each of the class are discussed late except for late stage ER/PR+/HER-2- (no significant cluster was obtained).

2. **Quantitative-real-time polymerase-chain-reaction**

**2.1 Cell culture**

The MCF-7 cells were cultured in Dulbecco’s Modified Eagle’s Medium (DMEM) whereas T47D and MDA-MB-231 cells in Roswell Park Memorial Institute medium (RPMI) supplemented with 10% fetal bovine serum (FBS) and penicillin-streptomycin (MP Biomedicals) at 37 °C, 5% CO2 and 95% humidity. MCF10A, a kind gift from Dr. Annapoorni Rangarajan (IISC, Bangalore, India) was maintained in DMEM F12 containing horse serum supplemented with hydrocortisone, EGF, insulin, cholera toxin and penicillin-streptomycin at 37 °C, 5% CO2 and 95% humidity. The cells were grown until 70-80% confluence and then sub-cultured with Trypsin-EDTA.

**2.2 Total RNA isolation and qRT-PCR**

Total RNA was isolated from MCF7, T47D, MDA MB-231 and MCF10A cells using Tri-reagent (Sigma-Aldrich). A total of 500ng was digested with DNase-I enzyme (Sigma-Aldrich) and was subjected to cDNA synthesis using superscript II first strand synthesis kit (Thermo-Scientific). Reverse transcription PCR and qRT-PCR was performed using primers. The primer sequences were as follows: MAML2-forward 5’- CAAATGCAGAATCAGCCCATTGCA-3’, reverse-5’- AGAAGAAGTTGCTGTTTCTGCTCC-3’, NLGN3-forward 5’- ATATTGCCTTCTTCGGGGGAGAC-3’, reverse-5’- AGACAGTCCACCATATCCACGGT-3’, TTN-forward- GCCAACAGTGGACGATATTCCCT, reverse-5’- AATCAGTAAGCTGTAGAGGTCGCC-3’, SYNE1-forward-5’ GTGATGCAGAGGCTGCAAGATGAG-3’, reverse-5’- GGGTCGGCCATCAGCTATATCG-3’, ANK2-forward-5’- TTGCTAAGCGTCTGGGCTACATC-3’, reverse-5’- CCTCCAAGCTGTATCGCAGGTAA-3’.

**3. Western blot analysis**

The RIPA buffer used for preparation of lysate was composed of 20 mM Tris-HCl (pH 7.5), 150 mM NaCl, 1 mM Na2EDTA, 1 mM EGTA, 1% NP-40, 1% sodium deoxycholate, 2.5 mM sodium pyrophosphate, 1 mM β-glycerophosphate, 1 mM Na3VO4 and 1 μg/mL leupeptin. The lysed samples were collected after centrifugation for 15 min at 12,000 rpm, 4 °C. Equal amount (30 μg) of proteins were loaded after Bradford method of protein quantification. The samples were run in 10% SDS-PAGE gel, transferred on PVDF membrane (Millipore) and blocked with 5% (w/v) non-fat milk (Sigma). Blots were then incubated with primary antibody overnight [MAML2 (1:5000) (ABCAM ab218338), ANK2 (1:5000) (SIGMA HPA008007), SYNE1 (1:5000) (SIGMA HPA019113), α-tubulin (1:1000), NLGN3 (1:1000) (SIGMA SAB5201608), Thereafter, 1h with their respective HRP conjugated secondary antibody [anti-rabbit (1:5000, Sigma Aldrich) or anti-mouse (1:5000, Sigma Aldrich)]. The protein bands were detected by using Gel Doc™ XR + Imager and subjected to densitometry analyses.
