## Supplementary tables and figures for "Stagewise identification of significantly mutated and dysregulated gene clusters in major molecular classes of breast invasive carcinoma": Supplementary_fig.S2c.pdf

### Regulation of the Epithelial Mesenchymal Transition pathway

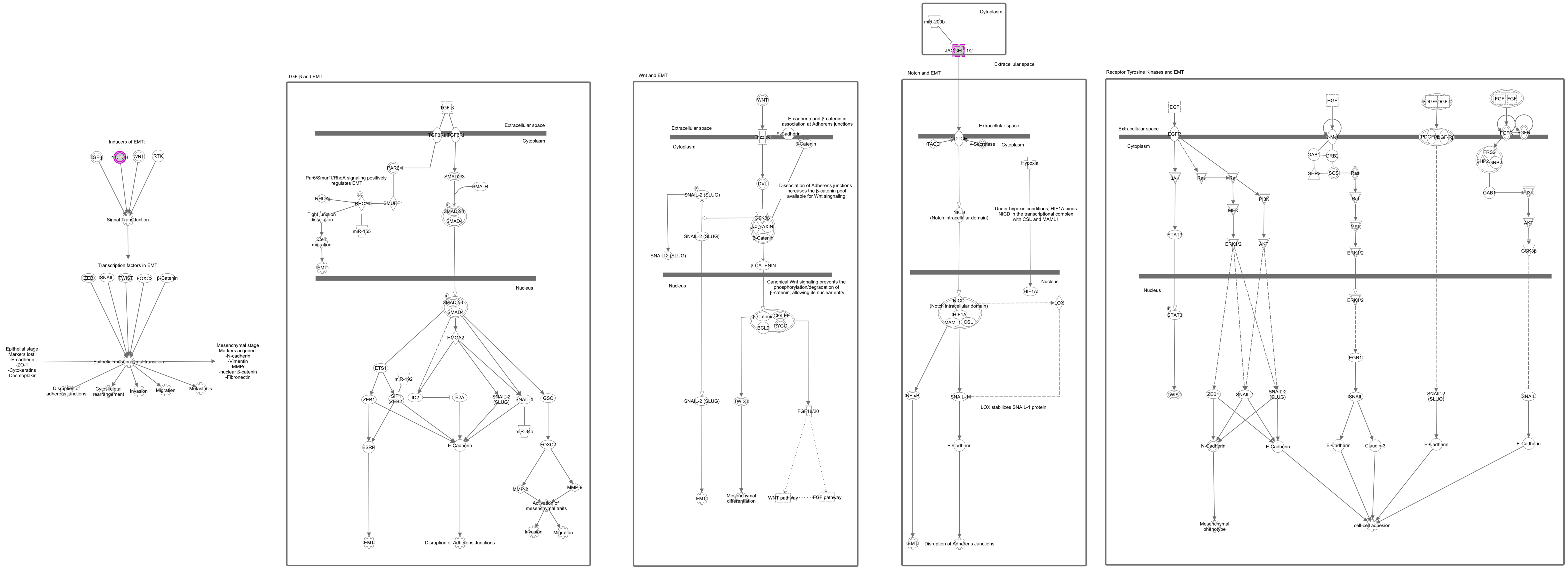


© 2000-2019 QIAGEN. All rights reserved.
