## Supplementary tables and figures for "Stagewise identification of significantly mutated and dysregulated gene clusters in major molecular classes of breast invasive carcinoma": Supplementary_fig.S4a.pdf

### Galactose degradation I(Leloir pathway)

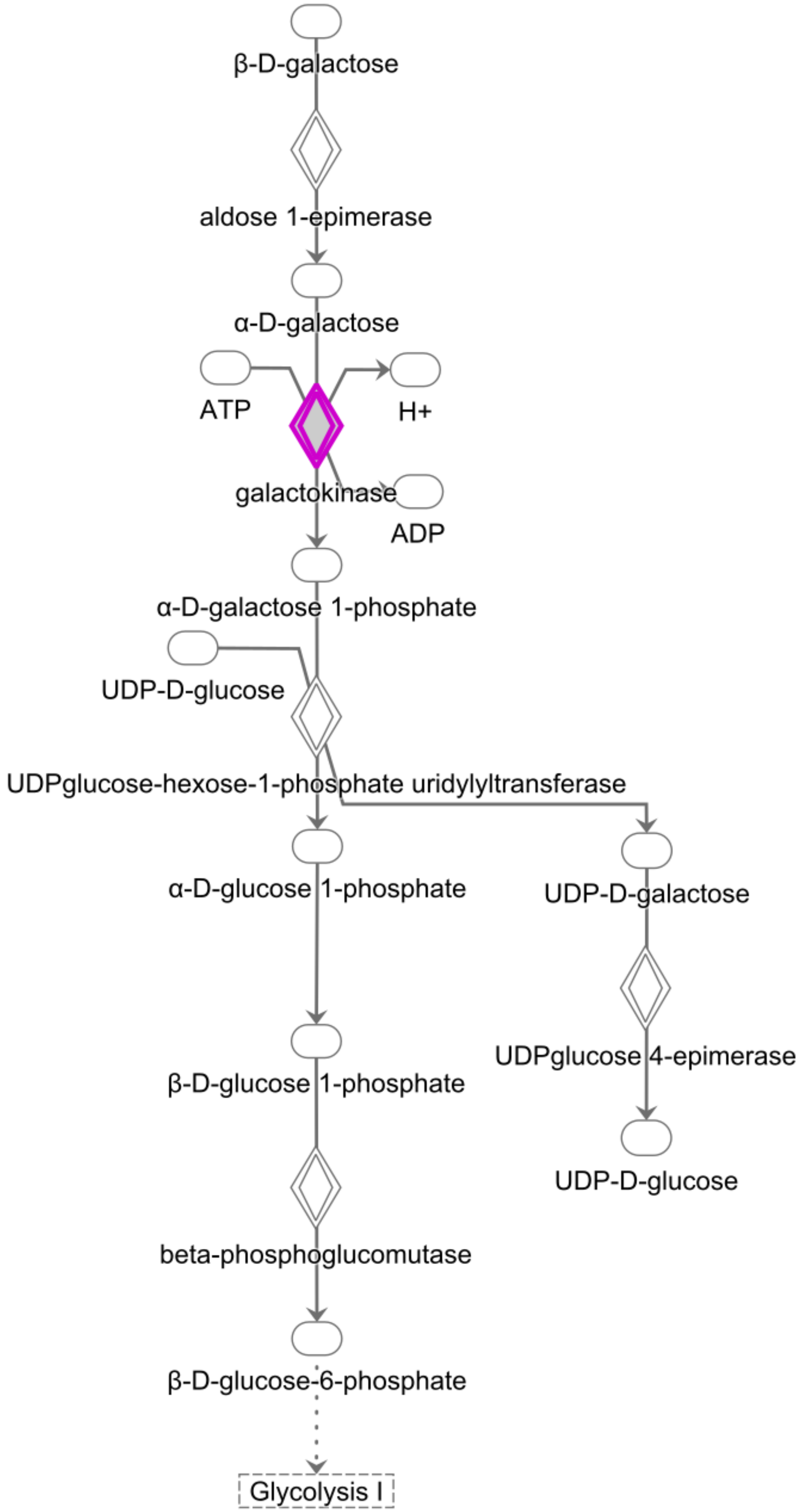


Adapted from HumanCyc.© 2000-2019 QIAGEN. All rights reserved.
