## Supplementary tables and figures for "Stagewise identification of significantly mutated and dysregulated gene clusters in major molecular classes of breast invasive carcinoma": Supplementary_fig.S4b.pdf

### Colanic Acid Building Blocks Biosynthesis

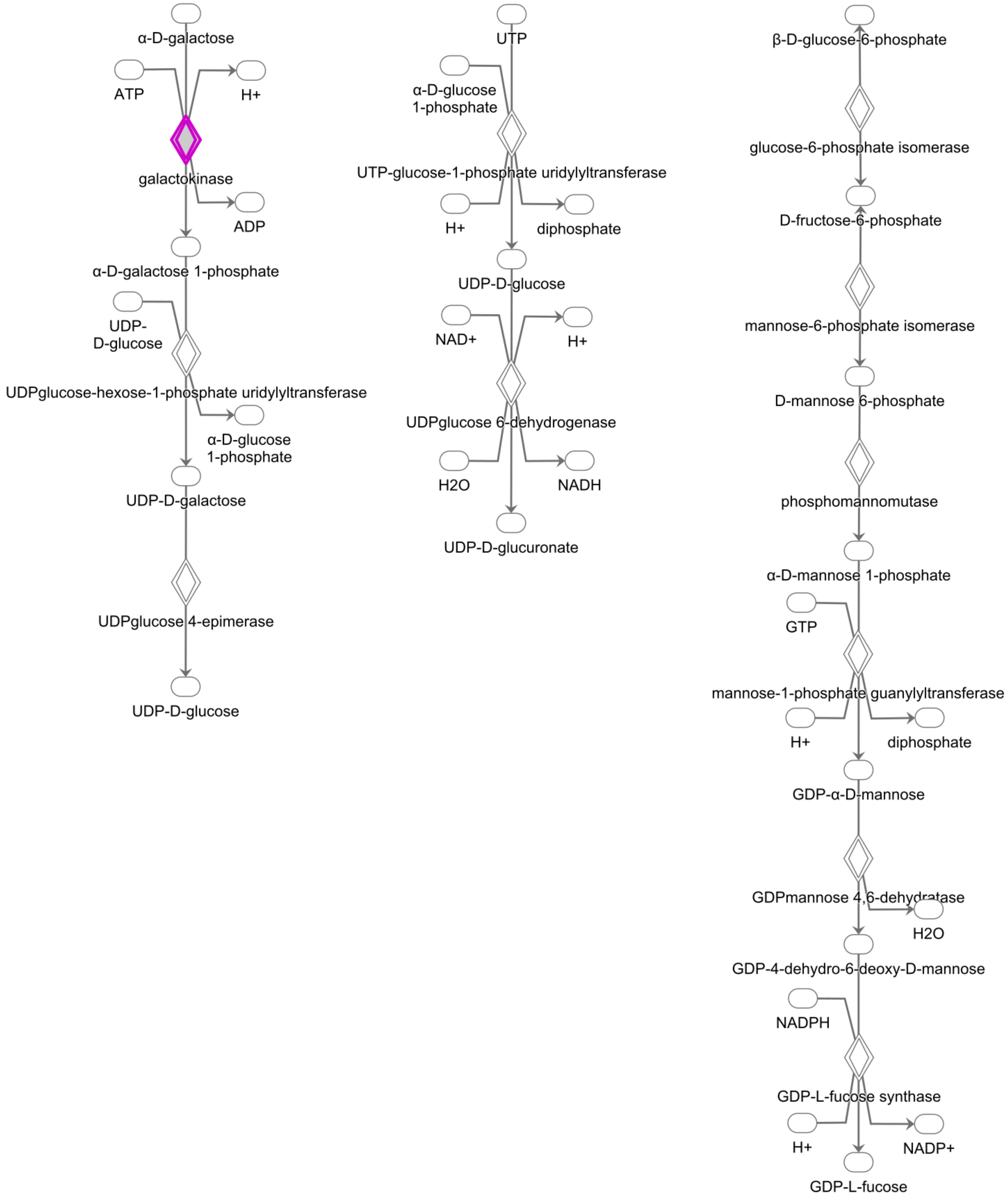


Adapted from HumanCyc.© 2000-2019 QIAGEN. All rights reserved.
