## Supplementary tables and figures for "Stagewise identification of significantly mutated and dysregulated gene clusters in major molecular classes of breast invasive carcinoma": Supplementary_figures_captions.docx

**Supplementary Figure S1. Figure showing pathway diagram of TR/RXR activation pathway mediated by NLGN3, NRXN1, NLGN1 network and found to be involved in early stage ER/PR+/HER-2- class of breast invasive carcinoma with p-value= 3.43e-03.**

**Supplementary Figure S2a. Figure showing pathway diagram of NOTCH signaling pathway mediated by NOTCH genes and interactors and found to be involved in early stage ER/PR-/HER-2+ class of breast invasive carcinoma with p-value 2.88e-17.**

**Supplementary Figure S2b. Figure showing pathway diagram of Th1 and Th2 activation pathway mediated by NOTCH genes and interactors and found to be involved in early stage ER/PR-/HER-2+ class of breast invasive carcinoma with p-value 1.00e-07.**

**Supplementary Figure S2c. Figure showing pathway diagram of Epithelial mesenchymal transition (EMT) pathway mediated by NOTCH genes and interactors and found to be involved in early stage ER/PR-/HER-2+ class of breast invasive carcinoma with p-value 1.82e-05.**

**Supplementary Figure S3a. Figure showing pathway diagram of netrin signalling mediated by TTN and interactors and found to be involved in early stage TNBC with p-value 2.61e-02.**

**Supplementary Figure S3b. Figure showing pathway diagram of RhoA signalling mediated by TTN and interactors and found to be involved in early stage TNBC with p-value 4.85e-02.**

**Supplementary Figure S3c. Figure showing pathway diagram of Axonal guidance signalling mediated by TTN and interactors and found to be involved in early stage TNBC with p-value 1.33e-02.**

**Supplementary Figure S4a. Figure showing pathway diagram of Galactose degradation I pathway mediated by DMD, ANK1, ANK2 and interactors and found to be involved in late stage TNBC with p-value 3.55e-03.**

**Supplementary Figure S4b. Figure showing pathway diagram of Colanic acid pathway mediated by DMD, ANK1, ANK2 and interactors and found to be involved in late stage TNBC with p-value 9.91e-03.**

**Supplementary Figure S4c. Figure showing pathway diagram of nNOS signaling pathway mediated by DMD, ANK1, ANK2 and interactors and found to be involved in late stage TNBC with p-value 2.81e-02.**

**Supplementary figure S5. Figure showing differential expression of genes a) NLGN3, b) MAML2 c) TTN d) SYNE1 and e) ANK2 from UALCAN. The x-axis represents normal and tumor samples and y-axis represents transcript per million values. All genes are downregulated in tumor samples as compared to normal samples.**

**Supplementary figure S6. Survival analysis of a) NLGN3, b) MAML2, c) SYNE1, d) TTN, e) ANK2, f) BRCA1 and g) BRCA2. The curve is plotted using the overall survival of cases (x-axis) with and without alteration in gene vs. the survival in months (y-axis). The curve represents that the overall survival of cases with alteration in NLGN3, MAML2, SYNE1 and TTN gene is less than the cases without alteration in genes.**
