## Supplementary figures and images for "Stagewise identification of significantly mutated and dysregulated gene clusters in major molecular classes of breast invasive carcinoma"

### Supplementary_fig.S1.pdf

Supplementary fig. S1

#### TR/RXR Activation pathway

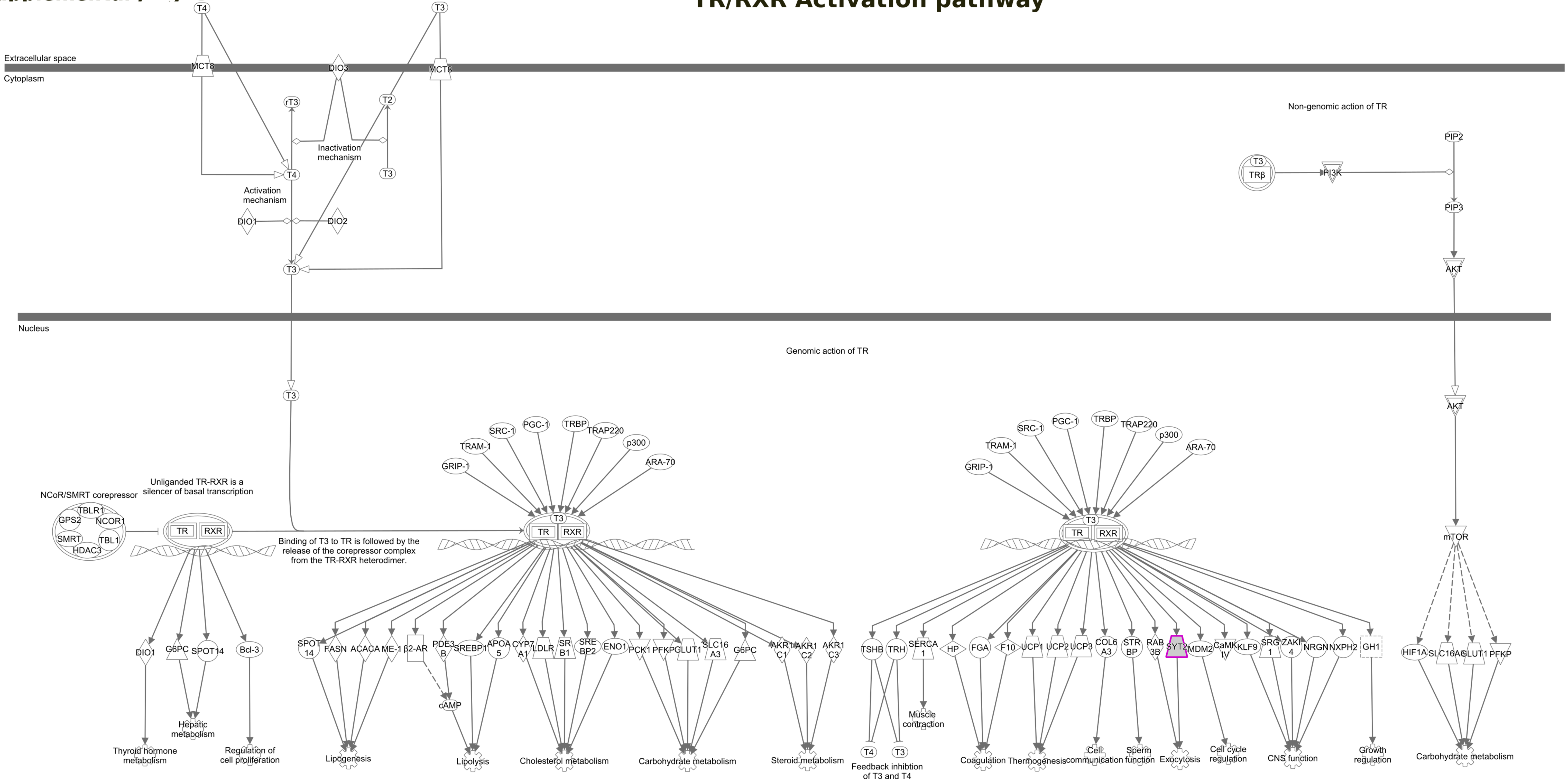


© 2000-2019 QIAGEN. All rights reserved.

### Supplementary_fig.S2a.pdf

Supplementary fig. 2a

#### NOTCH Signaling

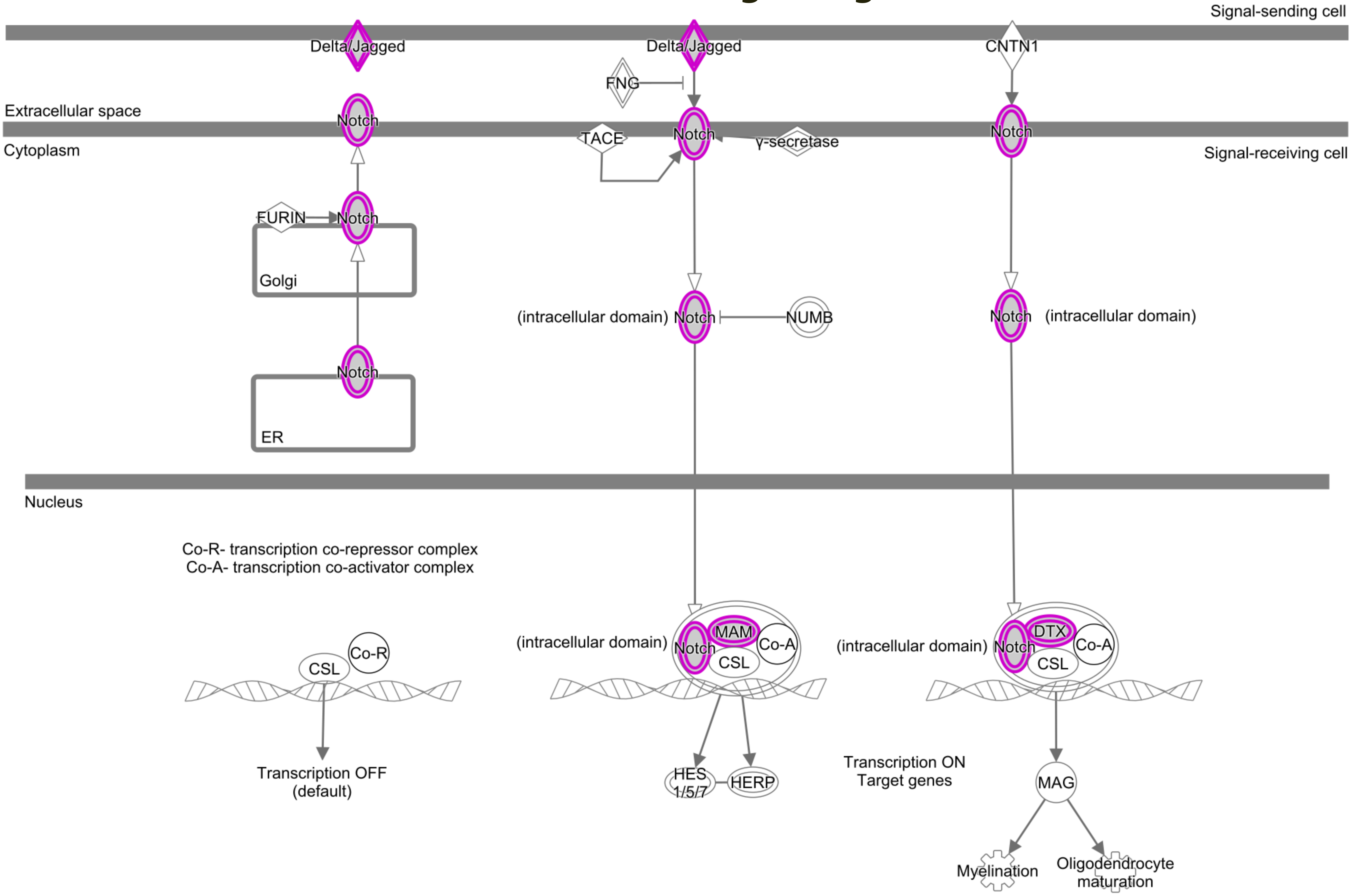


© 2000-2019 QIAGEN. All rights reserved.

### Supplementary_fig.S2b.pdf

Supplementary fig. S2b

#### TH1 and TH2 Activation pathway

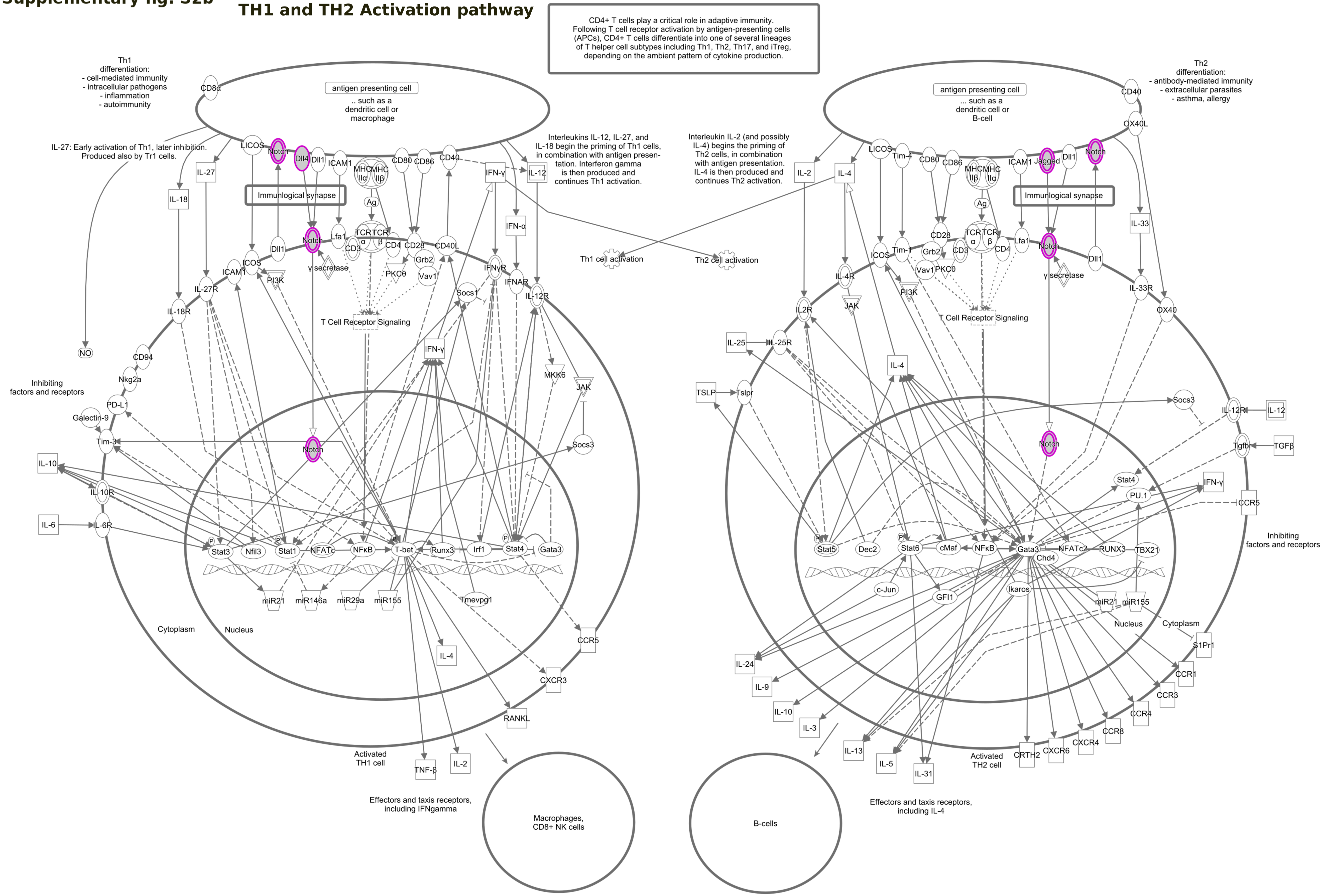


© 2000-2019 QIAGEN. All rights reserved.

### Supplementary_fig.S3a.pdf

Supplementary fig. 3a

#### Netrin Signaling

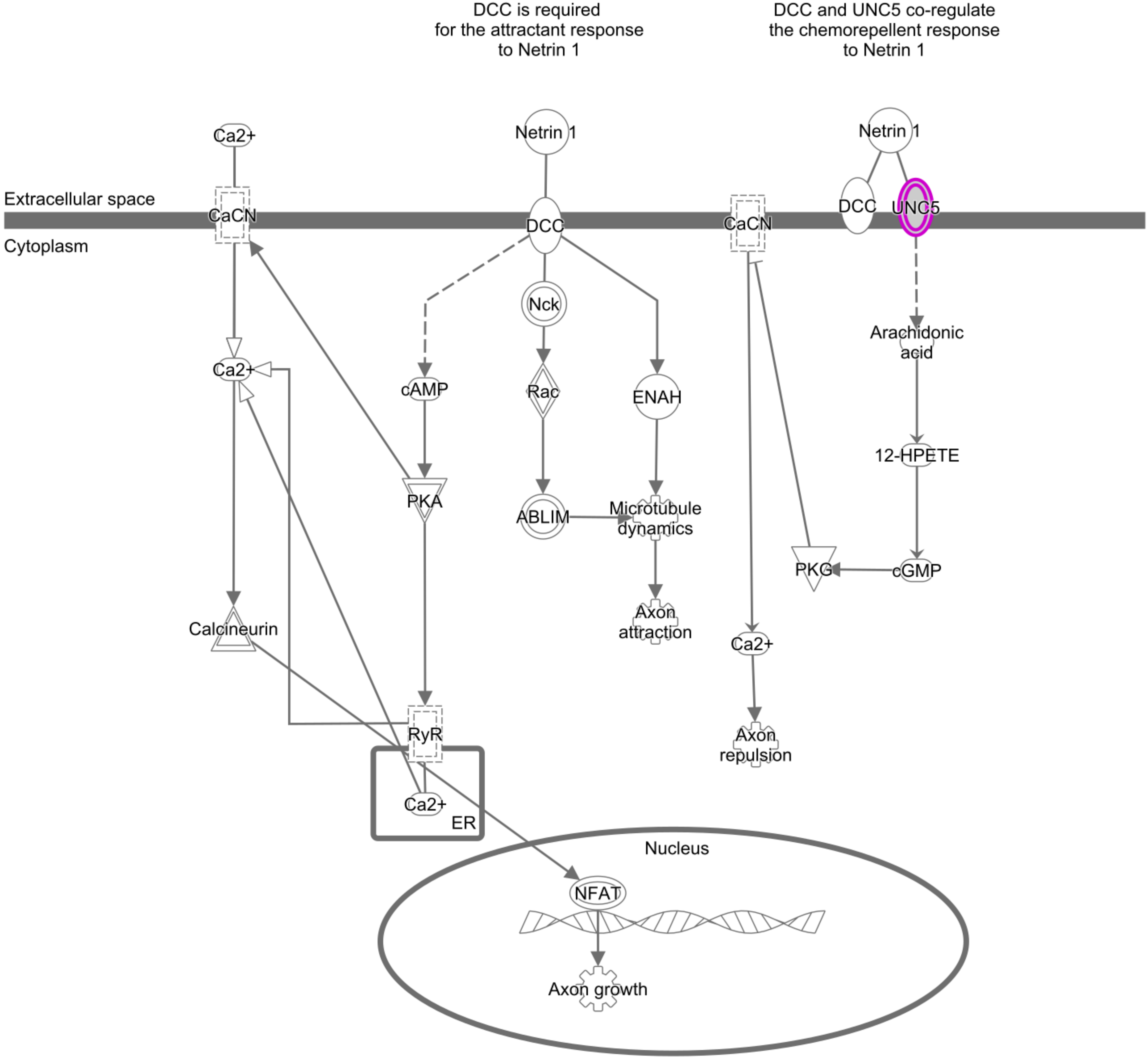


© 2000-2019 QIAGEN. All rights reserved.

### Supplementary_fig.S3b.pdf

Supplementary fig. S3b

#### RhoA signaling

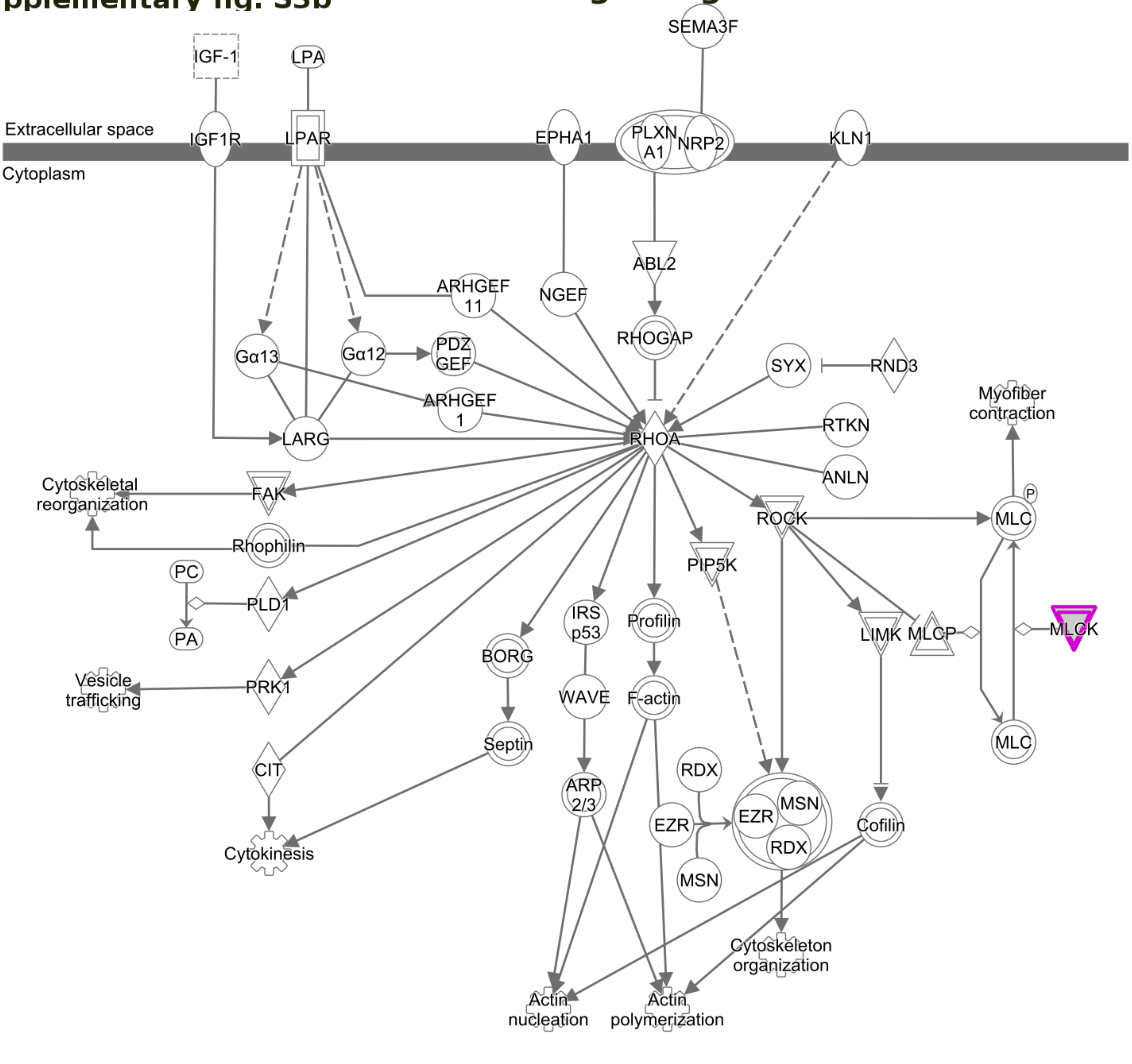


© 2000-2019 QIAGEN. All rights reserved.

### Supplementary_fig.S3c.pdf

**Supplementary fig. S3c**

#### Axonal guidance signaling

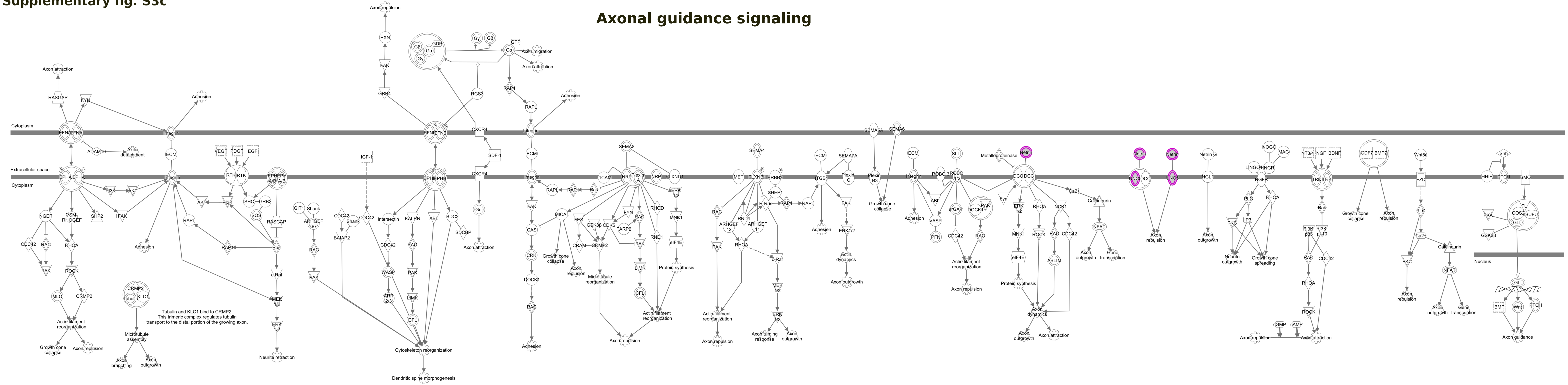


© 2000-2019 QIAGEN. All rights reserved.

### Supplementary_fig.S4c.pdf

**Supplementary fig. S4c**

##### nNOS signaling in skeletal muscle cells

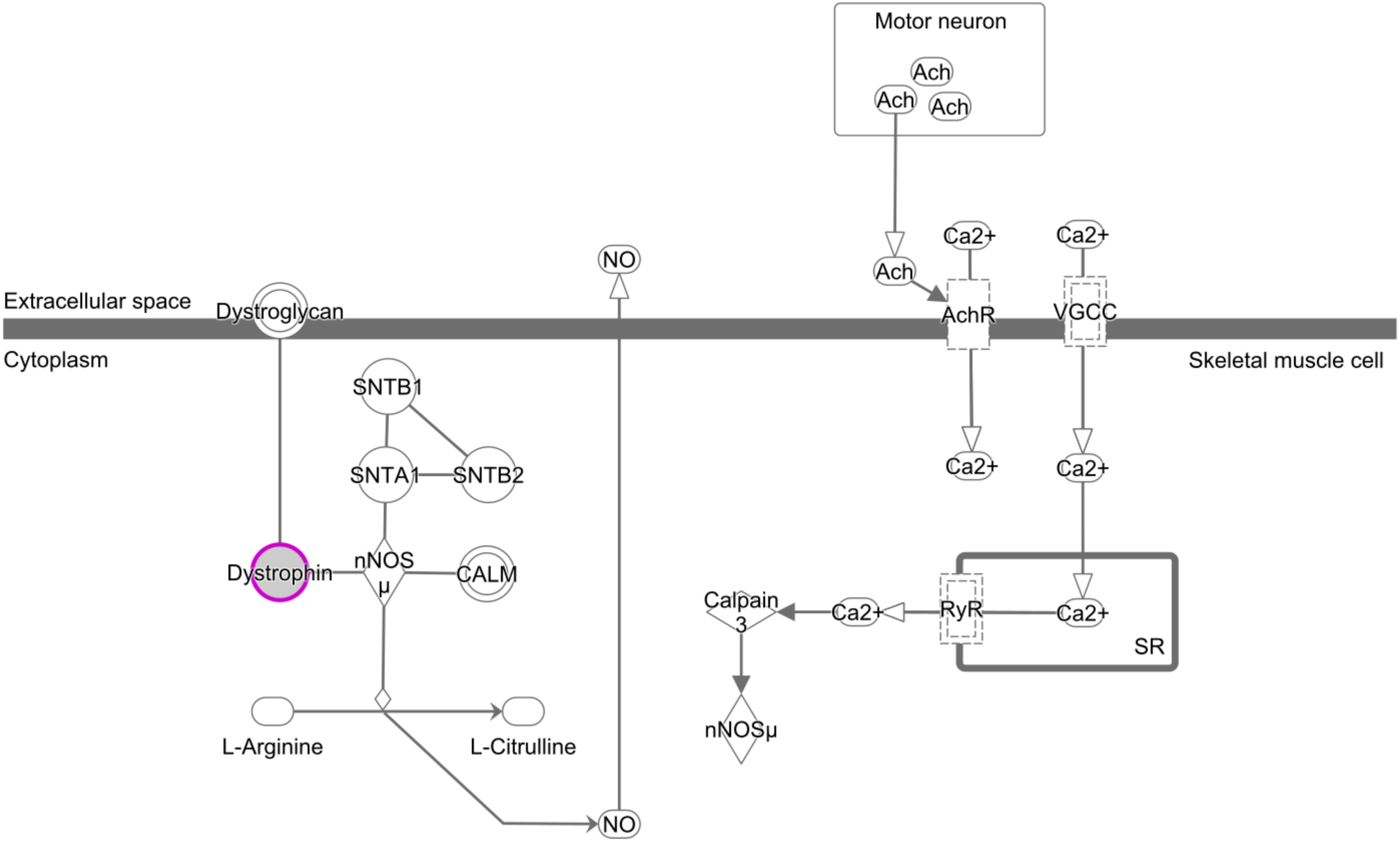


© 2000-2019 QIAGEN. All rights reserved.

### Supplementary_fig.S5.pdf

#### Supplementary fig. S5

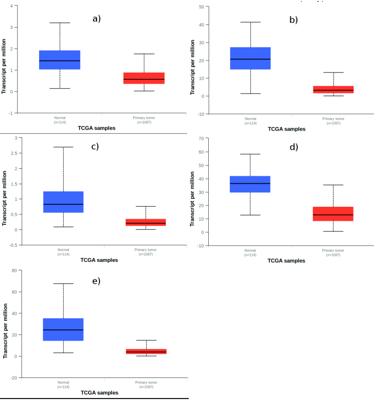

### Supplementary_fig.S6.pdf

##### Supplementary fig. S6

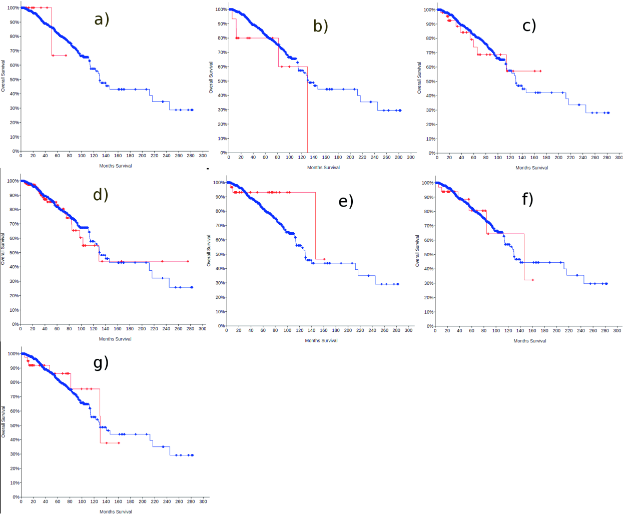
